# An Itch Receptor Drives Melanoma

**DOI:** 10.64898/2026.06.01.729383

**Authors:** Naina Gour, Aishwarya Atakkatan, Moloud Akbarzadeh, Hwan Mee Yong, Aishwarya Magesh, Zhiping Wu, Hyunkeun Joo, Yash Chhabra, Junmin Peng, Ashani Weeraratna, Mathieu Laplante, Stephane Lajoie, Xinzhong Dong

**Affiliations:** Solomon H. Snyder Department of Neuroscience, Howard Hughes Medical Institute, Johns Hopkins School of Medicine, Baltimore, MD, USA; Department of Otolaryngology, Head and Neck Surgery, Johns Hopkins School of Medicine, Baltimore, MD, USA; Department of Molecular Microbiology and Immunology, Johns Hopkins Bloomberg School of Public Health, Baltimore, MD, USA; Centre de recherche de l’Institut Universitaire de Cardiologie et de Pneumologie de Québec - Université Laval (IUCPQ-ULaval), Université Laval, Québec, QC, Canada; Department of Environmental Health and Engineering, Johns Hopkins Bloomberg School of Public Health, Baltimore, MD, USA; Departments of Structural Biology and Developmental Neurobiology, St. Jude Children’s Research Hospital, Memphis, TN, USA; Department of Biochemistry and Molecular Biology, Johns Hopkins Bloomberg School of Public Health, Baltimore, MD, USA; Cancer Signaling and Microenvironment, Fox Chase Cancer Center, Philadelphia, PA 19111, USA; Kompass Diagnostics, Chicago, IL, USA

## Abstract

Mas-Related GPCR X4 (MRGPRX4) is a sensory neuron-restricted receptor for cholestatic itch. Here, we identify MRGPRX4 as an unexpected melanoma driver. MRGPRX4 is upregulated in melanoma and is confined to invasive, neural-crest-like and pre-EMT states associated with dedifferentiation and therapy resistance. Ectopic MRGPRX4 expression in mouse melanocytes drives 100% penetrant, highly metastatic melanoma, demonstrating oncogenic behavior. MRGPRX4 promotes melanoma cell proliferation and invasion through basal, ligand-independent, PI3K-AKT-MAPK activation. Multi-omics links MRGPRX4 expression to a mesenchymal/neural-crest–like program that defines a distinct invasive MRGPRX4□ niche. Comparing transcptomics of MRGPRX4-driven tumors shows a broad overlap with BRAF/NRAS-driven tumors; however, the MRGPRX4 model also enriches an ECM-rich, invasive neural crest–like melanoma state. MRGPRX4 further remodels the tumor microenvironment toward an immunosuppressive, checkpoint-high state enriched in suppressive myeloid cells. Pharmacologic MRGPRX4 inhibition suppresses basal signaling and limits melanoma growth and invasion. In sum, melanoma exploits MRGPRX4 to acquire an invasive and immunosuppressive phenotype, nominating this somatosensory GPCR as a promising therapeutic target.

## Introduction

Conventional models of tumorigenesis emphasize genetic mutations that activate oncogenes or disable tumor suppressors. G protein-coupled receptor (GPCR) pathways fit this paradigm: mutations in GPCRs or their G-protein subunits occur in ∼20% of cancers and are widely viewed as the principal route to aberrant signaling^1^. However, GPCR oncogenicity is not restricted to coding mutations. Across solid tumors, GPCRs are frequently differentially expressed, their expression signatures correlate with oncogenic pathways and survival, and these programs can be largely independent of canonical driver mutations^2^.

Classic studies have shown that simple overexpression of unmutated GPCRs - and in some cases without ligand - can drive malignant transformation. This concept was first appreciated in the 1980s, when ectopic expression of the angiotensin receptor MAS1^3^ transformed NIH-3T3 fibroblasts in vitro and induced tumors in nude mice^4^. In line with this, aberrant expression of the neuronal serotonin receptor 5-HT1C/5-HT2C transformed NIH-3T3 fibroblasts and generated tumors in nude mice^3-5^. Overexpression of the normally neuronal metabotropic glutamate receptor GRM1/mGluR1 in melanocytes induced glutamate-dependent spontaneous melanoma in mice^6^. GPCRs can also transform in ligand-independent fashion, based on their high basal activities. GPR18, an orphan GPCR identified as an overexpressed orphan receptor in melanoma metastases and reported to be constitutively active^7^. Constitutive activity has also been described for other transformation-associated GPCRs, including 5-HT2C^8^, whose ectopic expression transforms NIH-3T3 cells and generates tumors in nude mice^5,8^. Together, these studies support a model in which aberrantly expressed wild-type GPCRs can promote relevant oncogenic programs through either ligand-amplified signaling or ligand-independent basal activity.

The cloning of MAS1 ultimately led to the identification of the Mas-related GPCR (MRGPR) family, defined by sequence homology to MAS1^9^. In primates, this family comprises multiple receptors, most of which are normally restricted to peripheral sensory neurons. The human MRGPRX subfamily contains four members: MRGPRX1, MRGPRX2, MRGPRX3, and MRGPRX4^10^. MRGPRX1 contributes to nociception^11^, MRGPRX2 mediates mast-cell pseudoallergic drug responses^12^, MRGPRX3’s function remains to be identified. We and others have shown that MRGPRX4 functions as an itch receptor in a small subset of sensory neurons^13-15^. To date, MRGPRXs have not been implicated in driving cancer, and no oncogenic functions have been described. Here, we show that MRGPRX4 - normally confined to a subpopulation of nociceptors behaves like a melanoma oncogene when aberrantly expressed in melanocytes, which share common progenitors (neural crest cells) with nociceptors. These findings identify MRGPRX4 as a previously unrecognized melanoma-associated oncogenic GPCR and define a model of tumorigenesis rooted not in receptor mutation, but in lineage-inappropriate receptor expression and ligand-independent basal GPCR signaling.

## Results

### MRGPRX4 is highly expressed in human melanoma

Comparison of MRGPRX4 expression from normal skin (from Genotype-Tissue Expression; GTEx) and skin cutaneous melanoma (SKCM) samples from The Cancer Genome Atlas (TCGA) showed a tumor-specific upregulation of *MRGPRX4* mRNA (**Fig. 1A**, p=1.5x10^-145^). In line with this, we also find high expression of MRGPRX4 in uveal melanoma as compared to all other non-melanoma in TCGA cancers (**Fig. 1B**). MRGPRX4 is only known to be primarily expressed in a small subset of human dorsal root ganglion (DRG) sensory neurons^15^, we next ascertained whether *MRGPRX4* could be expressed by human melanocytes. In bulk RNA-seq of primary human cultured skin melanocytes, *MRGPRX4* expression was undetectable despite robust expression of the canonical melanocyte markers *TYR* and *MITF* (**Fig. 1C**). Similarly, *MRGPRX4* was not detected in normal human skin keratinocytes (**Fig. 1D**) or fibroblasts (**Fig. 1E**), despite strong expression of the corresponding lineage markers *KRT14* and *VIM/COL1A2*, respectively. (**Fig. 1D, E**). Consistent with these findings, scRNA-seq analysis of normal human skin (**Supplementary Fig. 1A**) confirmed that *MRGPRX4* is not expressed by melanocytes, fibroblasts, keratinocytes, or eccrine tissue (**Supplementary Fig. 1B**).

**Figure 1.**
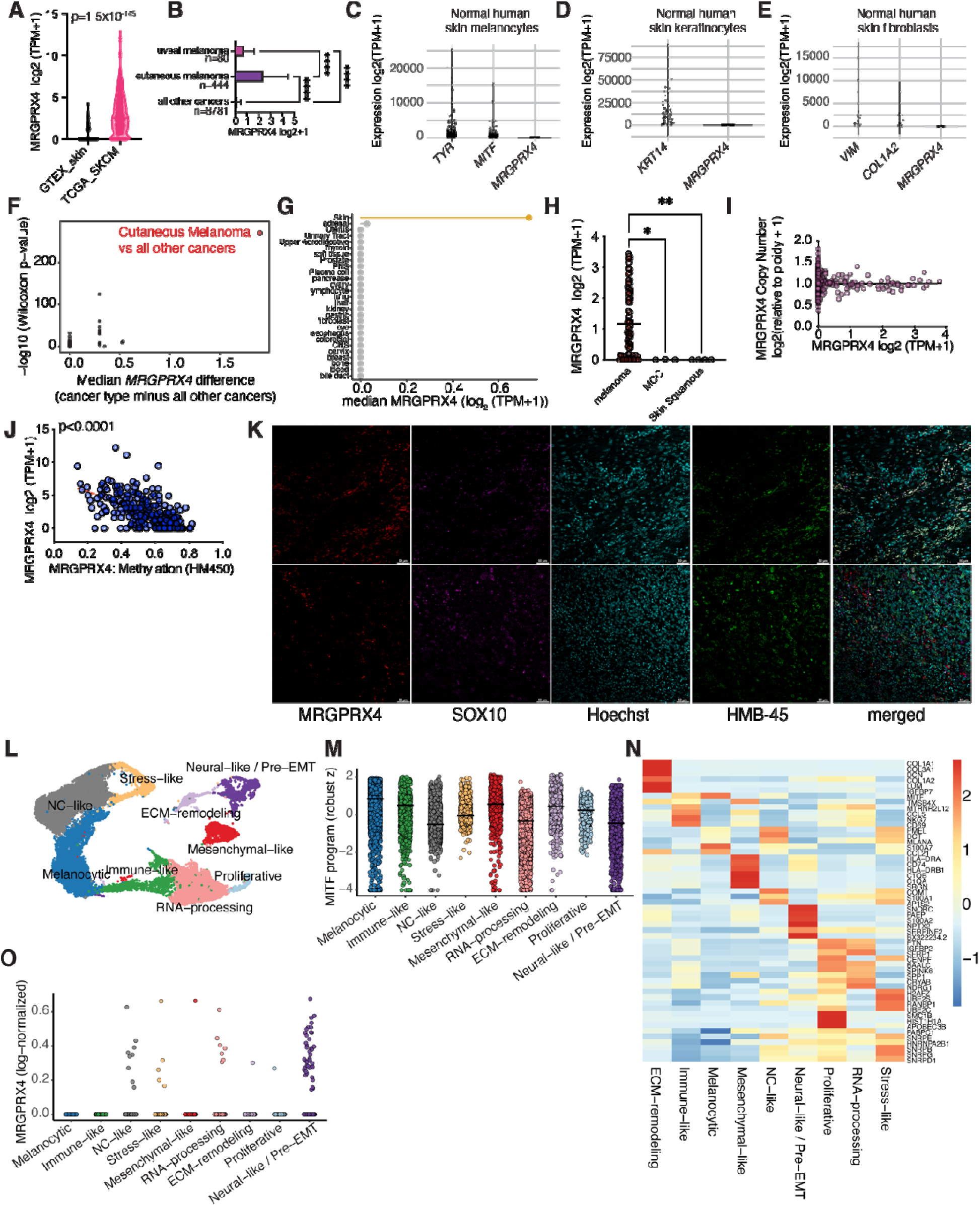
MRGPRX4 is highly expressed in human melanoma. (A,B) *MRGPRX4* mRNA expression in (A) GTEx normal skin and TCGA melanoma (SKCM) and(B) in uveal melanoma compared to other TCGA cancers. Data is analysed using Mann-Whitney U test or one-way ANOVA followed by post-hoc test. (C-E) *MRGPRX4* and canonical gene expression in normal human skin (C) melanocytes, (D) keratinocytes and (E) fibroblasts. (F) Cancer type-specific enrichment of *MRGPRX4* across TCGA PanCancer Atlas malignancies. Each point represents one TCGA cancer type. The x-axis shows the median *MRGPRX4* expression difference between that cancer type and all remaining TCGA cancers, and the y-axis shows the −log10 Wilcoxon p-value for the same one-vs-rest comparison. Cutaneous melanoma is highlighted in red and represents SKCM compared with all non-SKCM cancers. (G) Cell type-specific enrichment of *MRGPRX4* in DepMap cancer cell line data *(H) MRGPRX4* expression in DepMap skin cancer cell lines subdivided by cancer type. Data is analysed by one-way ANOVA followed by post-hoc test. (I) Relationship between *MRGPRX4* copy number and expression in TCGA SKCM patients (J) Relationship between *MRGPRX4* methylation and expression in TCGA SKCM patients. Red dotted line indicates best-fit linear regression (p<0.001). (K) Confocal images of human cutaneous melanoma sample stained for MRGPRX4 (red), SOX10 (magenta), HMB-45 (green) and Hoechst (cyan) nuclear counterstain. 20x magnification, scale bar 50um. (L) UMAP of integrated melanoma patient scRNA-seq datasets annotated into major melanoma transcriptional states. (M) MITF program scores across melanoma transcriptional states, showing highest activity in melanocytic cells and reduced activity in dedifferentiated lineages. (N) Heatmap of top cluster genes enriched in melanoma transcriptional states *(O) MRGPRX4* expression in melanoma transcriptional states

Next, we determined *MRGPRX4* expression levels across 29 TCGA cancer types. Cutaneous melanoma emerged as the malignancy with both the greatest increase in median *MRGPRX4* expression and the highest statistical significance (**Fig. 1F** and **Supplementary Fig. 1C**). Consistent with this, analysis of *MRGPRX4* expression in tumor cell lines from various tissue from the Cancer Cell Line Encyclopedia (CCLE) revealed that those derived from skin displayed the highest median levels of *MRGPRX4* expression (**Fig. 1G**). Upon subdividing skin-derived tumor lines into melanoma, Merkel cell carcinoma, and cutaneous squamous cell carcinoma cohorts, elevated *MRGPRX4* expression was observed exclusively in melanoma (**Fig. 1H**). Copy-number analysis indicated that *MRGPRX4* remained largely diploid, with no influence on transcript abundance (**Fig. 1I**). However, hypomethylation of the *MRGPRX4* locus tracked closely with transcript expression (**Fig. 1J**), and promoter methylation was significantly reduced in melanoma samples compared with benign nevi (**Supplementary Fig. 1D**). These findings suggest that epigenetic reprogramming contributes to aberrant *MRGPRX4* upregulation in melanoma. Lastly, we found that *MRGPRX4* mutations are very rare in melanoma. DepMap mutation data revealed damaging *MRGPRX4* variants in only 4 out of 1807 cancer cell lines (**Supplementary Fig. 1E**), and none among 148 melanoma cell lines (**Supplementary Fig. 1E**).

Analysis of the TCGA pan-cancer dataset showed a similar pattern, where only 8 of 468 SKCM tumors (1.7%) contained a protein-altering *MRGPRX4* mutation (**Supplementary Fig. 1F**), and no TCGA cancer type exceeded a mutation frequency of ∼4% (**Supplementary Fig. 1G**). These findings indicate that *MRGPRX4* is not a recurrently mutated driver in melanoma and that any observed alterations are likely rare, isolated events. Next, we performed multiplex immunofluorescence staining on primary human cutaneous melanoma sections from two patients. High-resolution tile-scans of large tumor areas showed that MRGPRX4 expression is restricted to discrete intratumoral areas rather than present across the entire lesion (**Supplementary Fig. 1H**). Tile-scanned sections were used to identify regions with enriched MRGPRX4 positivity, which were subsequently imaged at higher resolution together with SOX10 and HMB-45 (**Fig. 1K**). Across both tumors examined, MRGPRX4 cells were readily detected within regions strongly positive for SOX10 and HMB-45.

To first survey *MRGPRX4* expression within established melanoma transcriptional states, we analyzed DepMap melanoma cell lines using the framework identified by Tsoi *et al*.^16^, which stratifies melanomas along the melanocytic-mesenchymal axis into melanocytic, neural crest-like, transitory, and undifferentiated phenotypes (**Supplementary Fig. 1I**). PCA cleanly recapitulated this architecture, and overlaying *MRGPRX4* levels revealed its expression is predominantly localized to the neural-crest-like and transitory regions (**Supplementary Fig. 1J**). These states represent intermediate, invasive, and epithelial-mesenchymal transition (EMT)-leaning phenotypes rather than fully melanocytic or fully dedifferentiated melanomas. Consistent with this, re-analysis of the Tsoi *et al*. melanoma cell line panel (GSE80829) using their four-state classification showed that *MRGPRX4* mRNA expression peaks in neural crest-like and transitory lines and is lowest in undifferentiated lines (**Supplementary Fig. 1K**). These findings suggested that MRGPRX4 may be preferentially expressed in tumors occupying partial EMT/neural-like transcriptional states, prompting us to test this relationship at single-cell resolution in a dataset with finer melanoma state annotations. For this, we analyzed a publicly available dataset of human cutaneous melanoma (GSE269936). UMAP clustering of melanoma cells revealed several melanoma states (**Fig. 1L**), consistent with previously defined cellular programs^16,17^. MITF is a central determinant of melanoma cell differentiation (**Fig. 1M**). It drives the expression of genes involved in melanogenesis, the hallmark function of mature melanocytes. Malignant cells display a broad spectrum of MITF expression and activity levels (**Fig. 1M**). Distinct transcriptional states correlate with varying MITF activity, defining a melanocytic program characterized by high MITF activity and pre-EMT or neural crest-like (NC-like) states marked by low MITF activity (**Fig. 1M**). Signature gene expression for each melanoma state was then established (**Fig. 1N**). Using this melanoma state framework, we evaluated *MRGPRX4* expression. Here, we found that neural-like/pre-EMT cells are the predominant source of *MRGPRX4* expression (**Fig. 1O**). Together this data positions MRGPRX4 as a potential marker and effector of dedifferentiated, migratory programs associated with invasion and metastasis.

### MRGPRX4 drives metastatic melanoma in mouse model

In order to understand the role of MRGPRX4 in melanoma, we ectopically expressed MRGPRX4 in mouse melanocytes *in vivo*. For this, we crossed Tyrosinase (*Tyr)*^CreER^ animals, a melanocyte specific CreER line to a MRGPRX4-ROSA26-loxP-stop-loxP line (MRGPRX4^LSL^)^13^. Post tamoxifen injection, we found that Tyr^CreER+^;MRGPRX4^LSL+/-^ mice developed melanoma. This phenotype was 100% penetrant, seen in both males and females, and all experimental mice succumb to tumor burden around 3-5 months post tamoxifen injection. These animals exhibited widespread disease, including cutaneous (**Fig. 2A,B**), uveal (**Fig. 2C**), and meningeal melanomas (**Fig. 2D**). This was accompanied by an aggressive metastatic phenotype where draining lymph nodes were heavily infiltrated with melanoma cells (white arrows, **Fig. 2E**), and tumor nodules were also present in the lungs (white arrows, **Fig. 2F**) and brain parenchyma (**Fig. 2G**). While *Tyr*^CreER-^;*MRGPRX4*^LSL+/-^ controls (*Tyr*^CreER^-negative) lacked detectable expression, tumors from *Tyr^CreER+^*;*MRGPRX4*^LSL+/-^ mice confirmed robust *MRGPRX4* expression by immunofluorescence (**Fig. 2H**). Notably, tamoxifen-injected Cre-negative control mice remained tumor-free for up to 18 months (**Supplementary Fig. 2A**).

**Figure 2.**
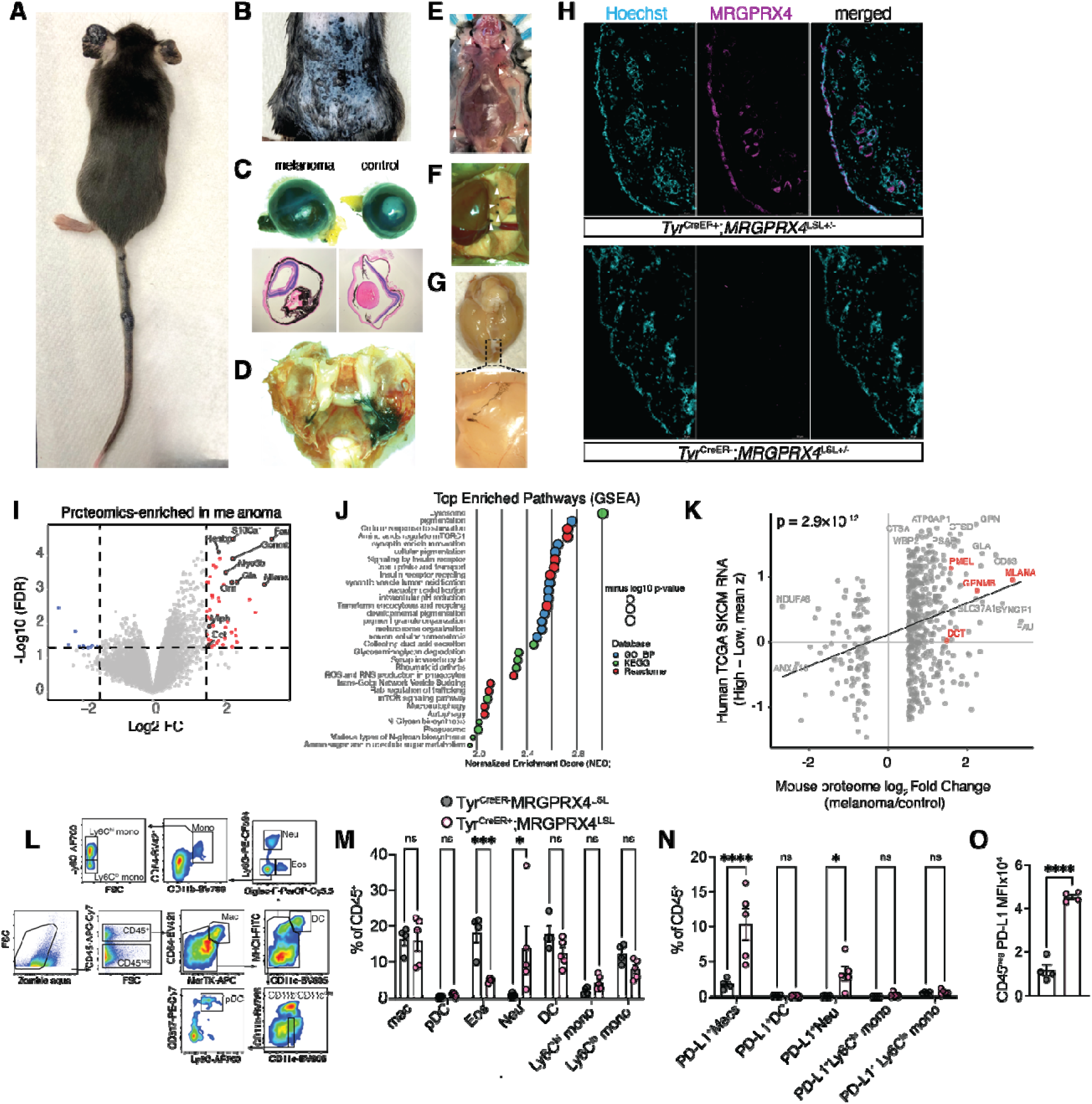
MRGPRX4 drives metastatic melanoma in mouse model. (A-G) (A) Representative image of melanoma on the ears and tails of tamoxifen-treated *Tyr*^CreER+^;*MRGPRX4*^LSL+/-^ mice (11 weeks old). (B) Cutaneous melanoma visualized following depilation. (C) Gross and H&E images of uveal melanoma from control *Tyr*^CreER-^;*MRGPRX4*^LSL+/-^ and melanoma *Tyr*^CreER+^;*MRGPRX4*^LSL+/-^ animals. (D) Meningeal melanoma visualized in *Tyr*^CreER+^;*MRGPRX4*^LSL+/-^ mouse after removal of skull and brain. *Tyr^CreER+^*;*MRGPRX4*^LSL+/-^ melanoma animals showing (E) lymph node metastasis (white arrows), (F) lung metastasis (white arrows) and (G) melanoma cells in brain parenchyma. (H) Tile-scan confocal images of MRGPRX4 staining in tamoxifen-treated experimental (*Tyr*^CreER+^;*MRGPRX4*^LSL+/-^) and control (*Tyr*^CreER-^;*MRGPRX4*^LSL+/-^) littermate mice at 20x magnification. Data is representative of 2-3 independent experiments. Scale bar, 50um (I-J) Proteomics analysis comparing tumor (*Tyr*^CreER+^;*MRGPRX4*^LSL+/^) vs control (*Tyr*^CreER-^ ;*MRGPRX4*^LSL+/-^) skin (I) Volcano plot where red dots are proteins significantly enriched in melanoma, and blue dots indicate significantly decreased proteins. (J) Top pathways enriched in melanoma skin samples (GSEA). Bubble plot shows normalized enrichment scores (NES), where bubble diameter represents statistical significance (-log10 p-value) and color indicates pathway database (GO biological process, KEGG, Reactome). (K) Cross-species conservation between MRGPRX4-driven melanoma proteome and human SKCM transcriptomics. Mouse protein log2 fold-change values (melanoma/control) aligned to orthologous human RNA expression (TCGA SKCM; z-scored). Genes enriched in mouse tumors also show elevated expression in human melanoma (p = 2.9 × 10 ¹², linear regression) (L-O) Immune microenvironment in MRGPRX4-driven melanoma. (L) Gating scheme for the identification of tumor myeloid celltypes. Frequencies of (M) myeloid cells and (N) PD-L1^+^ myeloid cells and (O) PD-L1 MFI of CD45^neg^ population in skin samples of control (*Tyr*^CreER-^ ;*MRGPRX4*^LSL+/-^) vs melanoma mice (*Tyr*^CreER+^;*MRGPRX4*^LSL+/-^). Data are mean+SEM and each dot represent an animal (n = 4-5), and are representative of 2-3 independent experiments. For all experiments both male and female mice were used and data are analyzed by Student’s t test or one-way ANOVA followed by post hoc test. *p < 0.05, **p < 0.01

Next, we confirmed if MRGPRX4-driven tumorigenesis was specific. For this, analogous genetic crosses with other MRGPRX cluster members using *MRGPRX1*^LSL^ ^18^, *MRGPRX2*^LSL^ ^19^, or *MRGPRX3*^LSL^ alleles with *Tyr*^CreER^ animals were generated. Unlike MRGPRX4, other family members i.e. MRGPRX1, MRGPRX2 and MRGPRX3 did not induce melanoma in vivo, (**Supplementary Fig. 2B**), indicating that this effect is specific to MRGPRX4.

To characterize MRGPRX4-driven melanoma, we performed mass-spectrometry-based proteomics analysis on cutaneous melanoma samples (*Tyr^CreER+^*;*MRGPRX4*^LSL+/-^) and skin from healthy control animals (*Tyr*^CreER-^;*MRGPRX4*^LSL+/-^). All samples showed similar overall intensity distributions between melanoma samples and control skin (**Supplementary Fig. 2C**) and robust genotype-associated proteomic differences when visualized on a PCA plot (**Supplementary Fig. 2D**). Unsupervised hierarchical clustering of the top differentially expressed proteins separated cutaneous melanoma samples from melanoma and control skin suggesting profound proteome-level changes (**Supplementary Fig. 2E**). Further analysis visualized via a volcano plot shows differentially abundant proteins in melanoma versus control skin (enriched proteins in tumor as red dots) including canonical melanocyte-lineage proteins (DCT, MLANA, GPNMB) (**Fig. 2I**). Gene Set Enrichment Analysis (GSEA) of melanoma-enriched proteins identifies top pathways related to pigmentation, melanosome organization, lysosome function, vesicle trafficking, and autophagy, consistent with active melanocytic and metabolic programs (**Fig. 2J**). Next, we wanted to compare MRGPRX4-driven melanoma in mice to human melanoma. To this end, we performed a cross-species comparison between the MRGPRX4-driven mouse melanoma proteome and the human TCGA SKCM transcriptome. We projected the mouse tumor proteome onto the melanoma transcriptome derived from the human TCGA SKCM dataset. MRGPRX4-driven tumors displayed a dominant mesenchymal/EMT-like invasion and RNA-processing/cell-cycle program. Gene colors indicate assignment to established TCGA melanoma state categories (**Supplementary Fig. 2F**). In line with this, we found a significant positive correlation (p = 2.9 × 10 ¹²) (**Fig. 2K**) and strong expression of canonical melanocyte-lineage markers (MLANA, GPNMB, DCT, PMEL) between our mouse model and human melanoma. This suggests that the proteins upregulated in our MRGPRX4-driven melanoma mouse model correspond to mRNAs elevated in human melanoma.

As the immune composition and phenotype of the tumor microenvironment is a major determinant of tumor evasion, we next aimed to evaluate the immune composition of MRGPRX4-driven melanoma. Flow-cytometric profiling revealed that MRGPRX4-driven melanoma (Tyr^CreER+^;MRGPRX4^LSL+/-^) exhibits a prominent reshaping of the tumor immune landscape compared with control animals (*Tyr*^CreER-^;*MRGPRX4*^LSL+/-^). We examined the myeloid compartment in these tumors (gating scheme, **Fig. 2L**) and found that eosinophils were reduced and neutrophils were upregulated (**Fig. 2M**). The granulocyte bias we observe in our model is similar to the neutrophil-high melanoma programs associated with poor outcomes in humans^20-27^. The frequency of macrophages, plasmacytoid DCs (pDCs), conventional dendritic cells and monocytes was unchanged in our model (**Fig. 2M**). However, we found that both PD-L1 macrophages and PD-L1 neutrophils were enriched (**Fig. 2N**), and PD-L1 expression (MFI) on CD45 cells increased several-fold (**Fig. 2O**), although it is unclear whether this is contributed by melanoma cells or the tumor stroma. The expansion of checkpoint-positive myeloid cells in our model suggests suppression of anti-tumor effector lymphocytes, consistent with MRGPRX4 signaling reprogramming the tumor-immune microenvironment toward a PD-L1^hi^, neutrophil-rich state - a phenotype linked to poor clinical response in melanoma patients.

### MRGPRX4-driven tumors display an enhanced NC/EMT signature

To understand the tumor microenvironment of MRGPRX4-driven melanoma, we performed spatial transcriptomic analysis on dorsal skin tumor samples from mice (n=3). Spatial gene expression data were obtained from tissue sections (**Fig. 3A**), which distinctly delineated both tumor and adjacent non-tumor regions. Notably, key anatomical compartments—including the epidermis, dermal stroma, dermal white adipose tissue (dWAT), underlying muscle, and melanoma were clearly resolved (**Fig. 3B,C**), enabling spatial mapping of transcriptional heterogeneity within the tumor microenvironment. To improve spatial resolution and more accurately resolve microenvironmental heterogeneity, we applied the BayesSpace algorithm^28^ to the Visium transcriptomic data. This approach enabled sub-spot resolution to improve the discrimination of cell populations. We used the AUCell algorithm^13^ to score individual spots for enrichment of transcriptional programs associated with melanocytic differentiation (**Fig. 3D**), mesenchymal transition (**Fig. 3E**), RNA processing (**Fig. 3F**), antigen presentation (**Fig. 3G**), and hypoxia (**Fig. 3H**). These gene signatures were derived from previously characterized melanoma states and tumor microenvironmental pathways^17^. Spatial enrichment maps reveal regions characterized by high expression of melanocytic markers, localized within the central tumor zone (**Fig. 3D**), while mesenchymal (**Fig. 3E**) and hypoxia-associated (**Fig. 3H**) programs are preferentially enriched at the tumor-stroma interface. Notably, antigen presentation (**Fig. 3G**) and RNA processing (**Fig. 3F**) signatures were distributed more diffusely across both tumor and adjacent stromal areas, consistent with immune cell infiltration and cellular stress responses. These data collectively support the presence of spatially organized heterogeneity within the MRGPRX4-driven melanoma microenvironment.

**Figure 3.**
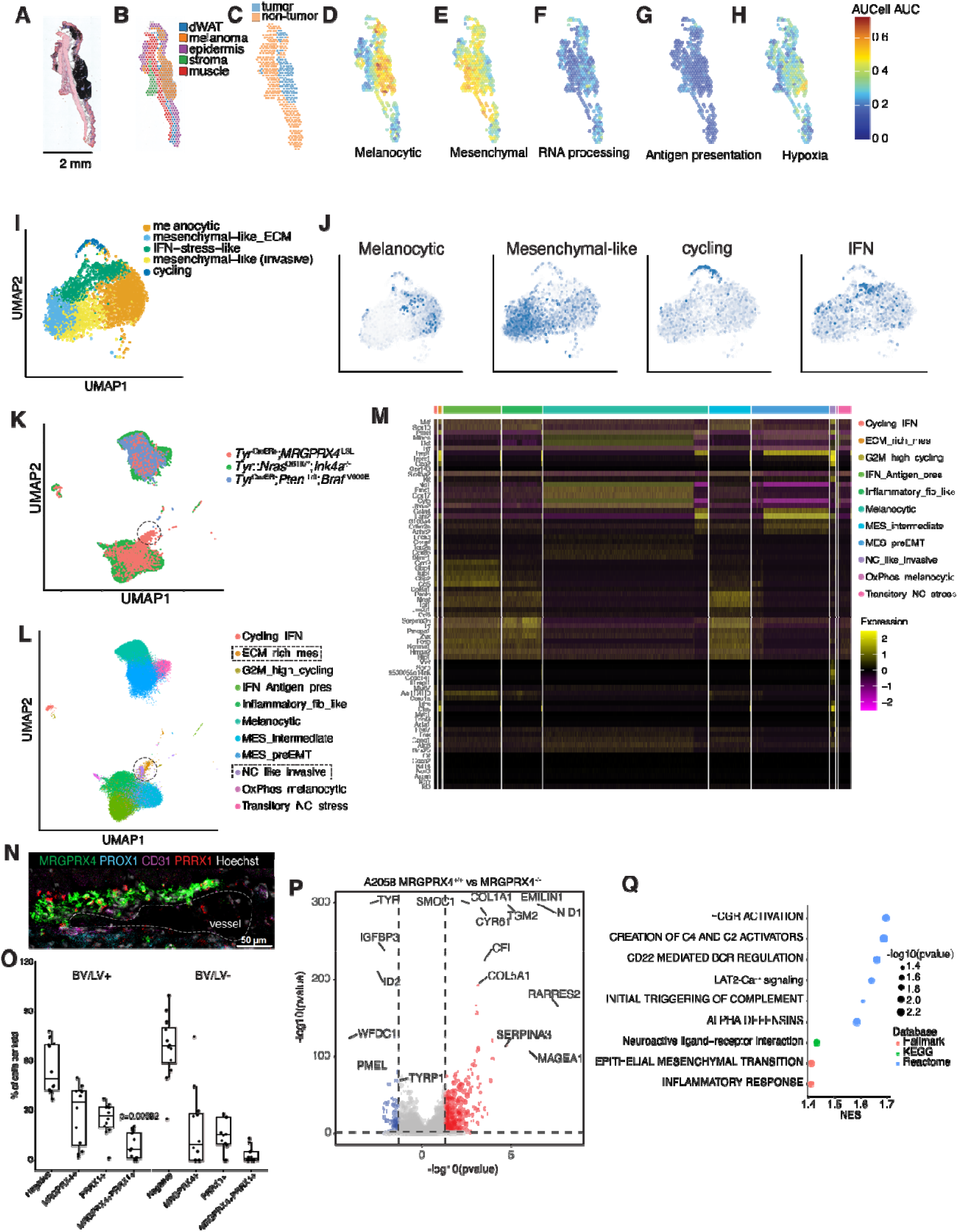
MRGPRX4 tumors display an enhanced NC/EMT signature. (A-H) Spatial transcriptomic of MRGPRX4-driven melanoma (*Tyr*^CreER+^;*MRGPRX4*^LSL+/-^). (A) Representative H&E-stained cross-section illustrating tumor, dermis, epidermis, stromal fascia,dWAT and muscle layers. Scale bar, 2 mm. (B) Spatial segmentation of Visium spots showing tissue compartment assignment based on canonical gene enrichment. (C) Spatial delineation of tumor-positive (blue) and tumor-negative (orange) regions (D–H) Spatial refinement using BayesSpace followed by AUCell scoring of predefined melanoma transcriptional programs. Shown are per-spot enrichment scores for (D) melanocytic, (E) mesenchymal-like, (F) RNA-processing, (G) antigen-presentation, and (H) hypoxia-associated states. Color intensity corresponds to AUCell score per spatial location. (I,J) (I) UMAP projection of dissociated melanoma cells colored by cell state, including melanocytic (orange), mesenchymal-like (hypoxia-associated; blue-green), IFN-stress-like (green), mesenchymal-like invasive (yellow), and cycling (dark blue) populations. (J) UCell-based scoring of melanoma-intrinsic programs shown at single-cell resolution of melanocytic, mesenchymal-like state, NCSC-like, and IFN-stress modules. Dot intensity reflects AUC score per cell. (K–M) (K) UMAP embedding of tumor-derived malignant cells colored by genotype, including *Tyr*^CreER+^;*MRGPRX4*^LSL^ (red), *Tyr::NRAS*^Q61K/°^;*Ink4a*^−/−^ (green), and *Tyr*^CreER^;*Pten*^fl/fl^;*Braf* ^V600E^ (blue) models. (L) UMAP colored by transcriptional states as indicated. (M) Heatmap of canonical state-defining genes across malignant clusters. (N, O) (O) Representative multiplex confocal image showing melanoma cells expressing MRGPRX4 (green) and PRRX1 (red) positioned directly adjacent to CD31 blood vessels (magenta) and PROX1 lymphatic endothelium (cyan). Nuclei are stained with Hoechst (white). Scale bar, 50 µm. (O) Quantification of phenotypically defined tumor subpopulations in blood vessel (BV) and lymphatic vessel (LV) positive areas (BV/LV^+^) versus BV/LV fields. BV/LV regions show significant enrichment of dual-positive MRGPRX4 PRRX1 invasive cells as compared to BV/LV^-^ areas (paired Wilcoxon test, p=0.0059). Dots represent independent imaged fields. (P, Q) (P) Volcano plot of differential expression between A2058 *MRGPRX4* ^/^ and *MRGPRX4* ^/^ cells highlighting down-regulation of melanocytic lineage markers (e.g., *TYR*, *TYRP1*, *PMEL*) and up-regulation of mesenchymal and extracellular-matrix genes (e.g., *COL1A1*, *COL5A1*, *RARRES2*, *SERPINA3*). (Q) GSEA enrichment analysis. Circle size denotes -log (p-value); color indicates database (Hallmark, KEGG, Reactome).

To characterize the tumor ecosystem surrounding MRGPRX4-driven melanoma, we generated a single-nucleus RNA-sequencing (snRNA-seq) dataset from matched tumor-bearing skin samples. Clustering revealed a diverse cellular landscape including melanoma cells, multiple fibroblast populations, vascular and lymphatic endothelial cells, keratinocytes, adipocytes, tissue-resident Schwann cells, immune infiltrates (including macrophage subtypes, T cells, and dendritic cells), and stromal populations (**Supplementary Fig. 3A**). These major compartments were transcriptionally distinct, as confirmed by cluster expression profiles (**Supplementary Fig. 3B**). Next, we isolated melanoma cells and performed downstream embedding and scoring exclusively within this compartment. This melanoma-only representation (**Fig. 3I**) revealed transcriptionally distinct melanoma states in line with others (**Fig. 3J**) ^16,17,29^. Using UCell scoring of curated melanoma gene sets, we identified five programs, melanocytic, two mesenchymal-like clusters, IFN-responsive, and cycling, which together span variation within our tumor model.

Next, we wanted to compare our model with well-established genetically engineered mouse models (GEMMs) of melanoma, including BRAF^30^ and NRAS^31^-driven models. To do this, we integrated our snRNA-seq data with publicly available data (GSE207592) from the *Tyr*::*NRAS*^Q61K/°^;*Ink4a*^−/−^ and *Tyr*^CreER+^;*BRAF*^V600E^;*Pten*^fl/fl^ models (**Fig. 3K**). Integrating these datasets revealed that several canonical melanoma states, including melanocytic, IFN-stress-like, mesenchymal-like, cycling, and antigen-presentation-associated populations, were shared across all three genotypes (**Fig. 3L**). In addition, MRGPRX4 tumors contained two divergent intermediate mesenchymal/neural crest-like populations that were minimally represented in the BRAF- and NRAS-driven tumors (**Fig. 3K–L**, dotted regions). Although these MRGPRX4-enriched compartments appeared as spatially distinct regions in the global integrated UMAP, they represented a substantial fraction of MRGPRX4 tumor cells. Together, these compartments comprised approximately half of the tumor-cell population in our model and were detected across all three MRGPRX4 tumors, representing 70.7%, 49.3%, and 42.5% of MRGPRX4 tumor cells in each sample. These compartments were also consistently enriched for invasive/intermediate melanoma states, indicating that they represent reproducible and substantial MRGPRX4-associated melanoma populations rather than sample-specific or minor outlier populations.

One MRGPRX4-enriched population exhibited a pronounced matrix-depositing phenotype, with high expression of collagens (*Col1a1*, *Col1a2*, *Col3a1*) and ECM-remodeling transcripts (*Mmp14*, *Timp3*, *Chsy3*) (**Fig. 3M**). This transcriptional program parallels the MES/undifferentiated “stromal/ECM” state described in human melanoma, characterized by collagen-rich, TGF-β–associated ECM networks linked to invasion and therapy resistance^16,32^. The second MRGPRX4-enriched melanoma cluster exhibited a neural crest-leaning potentially invasive program, marked by induction of neuronal, axon-guidance, and excitability-associated genes (e.g., *Nrg1*, *Epha4*, *Il1rapl1*, *Dclk2*, *Cacnb4*, *Kcna6*, *Grik2*, *Slc6a17*). This signature is consistent with melanoma phenotype-switching models, in which cells occupy intermediate neural crest-like/mesenchymal-like states associated with invasion. plasticity, and therapy adaptation^16,29,33,34^. Notably, NRG1/ERBB3 signaling has been shown in melanoma models to promote invasive behavior and support metastatic competence/colonization, providing a mechanistic precedent for linking Nrg1-high programs with invasive potential^35-37^. In addition, the cluster also contained a contractile-like module (*Tnnt3*, *Ttn*, *Sgcg*, *Dtna*), which have been previously reported as either mutated, upregulated in melanoma, driven by the Mitf transcription factor or induced in BRAF inhibitor-resistant melanoma cells^38-42^. This state showed partial retention but attenuation of melanocytic lineage identity, consistent with a dedifferentiated intermediate/neural crest-like transitory melanoma program along the differentiation continuum. Overall, the combined pattern of attenuated melanocytic lineage identity, paired with neural crest-leaning features and an intermediate/transitory character aligns with NC-like/transitory melanoma states described across human tumors and models, which sit between melanocytic and fully mesenchymal phenotypes and are associated with invasion and adaptive resistance^16,29,33^.

Single-cell transcriptomics of our MRGPRX4-driven melanoma model revealed substantial overlap with clusters identified in other GEMM of melanoma (*Tyr*::*NRAS*^Q61K/°^;*Ink4a*^−/−^ and *Tyr*^CreER+^;*BRAF*^V600E^;*Pten*^fl/fl^), but also identified populations specifically enriched in the MRGPRX4 context: an ECM+ subset and a NC-like subset with neuronal/axon-guidance features. This enrichment is noteworthy because recent lineage-tracing work has identified NC-like and related intermediate populations as cooperative drivers of invasion and metastatic colonization^17,43^, and these states also drive resistance to targeted therapy and tumor relapse^29,44^. Whether these MRGPRX4-enriched subpopulations causally drive the metastatic phenotype observed in our model remains to be evaluated.

To understand how these transcriptional states map back into tissue context, we performed multiplex confocal imaging. MRGPRX4-expressing melanoma cells formed dense clusters directly adjacent to CD31 vascular endothelium and near PROX1 lymphatic structures (**Fig. 3N**). This spatial pattern is consistent with prior evidence that melanoma contains a perivascular niche enriched for dedifferentiated, neural crest-like and tumorigenic cell populations that exhibit state plasticity and contribute to metastatic competence^17^. Within these vessel-proximal regions, a substantial fraction of MRGPRX4 cells co-expressed PRRX1, which is in line with work from others showing PRRX1^+^ melanoma cells have an EMT/mesenchymal-like transcriptional program associated with invasion^17^, although PRRX1 expression can exert context-dependent effects on metastatic outgrowth^45,46^. Across matched fields from the same lesions, vascular-adjacent (within 5 um) regions (BV/LV) displayed significantly elevated frequencies of double-positive MRGPRX4 PRRX1 cells relative to the non-vascular areas (BV/LV) (paired Wilcoxon p = 0.0059; **Fig. 3O**). Prior studies have demonstrated that lymphatic invasion and increased lymphatic vessel density are associated with dissemination and poorer prognosis in melanoma^47^, and that melanoma–lymphatic endothelial cell interactions actively promote metastatic behavior^48^. Together, these findings suggest that MRGPRX4 PRRX1 melanoma cells preferentially occupy vascular and lymphatic interfaces where invasive and dissemination-competent programs may be selectively reinforced.

To expand our findings in mouse models, we wanted to determine whether MRGPRX4 contributed to invasive transcriptional programs in human melanoma cells. To this end, we used A2058 melanoma cells, which constitutively express *MRGPRX4* (**Supplementary Fig. 3C**). We generated *MRGPRX4*^-/-^ A2058 cells and performed RNA-seq on *MRGPRX4*^+/+^ and *MRGPRX4*^-/-^ A2058 cells. CRISPR editing produced clonal disruption of the *MRGPRX4* locus, with both clones displaying a uniform indels (+2 and -7 bp) at the target site, as determined by ICE (Inference of CRISPR Edits) analysis (**Supplementary Fig. 3D-G**). These indels are predicted to induce a frameshift and premature termination, resulting in complete loss of functional *MRGPRX4*. Sequencing *MRGPRX4*^+/+^ and *MRGPRX4*^-/-^ A2058 cells, we found that MRGPRX4 expression was associated with reduced melanocytic differentiation gene expression (*TYR*, *TYRP1*, *PMEL*) and reciprocal induction of extracellular matrix–associated and inflammatory mediators, including *RARRES2*, *SERPINA3*, *COL1A1*, and *IGFBP3* (**Fig. 3P**). Gene-set enrichment analysis further revealed enrichment of complement-initiating pathways, Fcγ receptor–associated signaling, α-defensin modules, and LAT2-dependent Ca² pathways alongside EMT-associated signatures in MRGPRX4-expressing cells (**Fig. 3Q**). Complement activation and Fcγ receptor signaling have been implicated in shaping inflammatory, angiogenic, and metastatic tumor microenvironments in melanoma and other cancers^49,50^, and α-defensin expression has been reported at invasive tumor margins and in association with disease progression^51,52^. Collectively, these transcriptional features place MRGPRX4^+^ melanoma cells within an inflammatory, matrix-remodeling program consistent with invasive melanoma states. Consistent with the transcriptional shift observed upon *MRGPRX4* deletion in A2058 cells, analysis of DepMap melanoma lines stratified by endogenous MRGPRX4 expression revealed a similar overall pattern. Volcano plot of MRGPRX4^hi^ versus MRGPRX4^lo^ lines showed preferential enrichment of NC/EMT-linked transcripts like *NGFR*, *NES*, *MIA*, *SERPINA3*, *COL9A3* and *COL19A1* (**Supplementary Fig. 3H**). Consistent with our data comparing MRGPRX4^+/+^ with MRGPRX4^-/-^ A2058 cells, pathway enrichment GSEA in MRGPRX4^hi^ versus MRGPRX4^lo^ melanoma lines (DepMap) highlighted ECM-receptor interactions, focal adhesion, TGFβ signaling, inflammatory signaling, and PI3K-Akt pathways (**Supplementary Fig. 3I**). Plotting individual genes associated with these pathways revealed significant upregulation of *PRRX1*, *SERPINE2*, *CTHRC1*, *TNC*, *SPARC*, and *VIM* in MRGPRX4^hi^ melanoma cells (**Supplementary Fig. 3J**), consistent with a de-differentiating invasive program with MES-like/ECM-remodeling features. Concordantly, MRGPRX4 directly regulates MES/ECM-associated transcripts, as *MRGPRX4*^-/-^ A2058 cells show reduced *COL5A1*, *COL1A1*, *EMILIN1*, and *SERPINA3* mRNAs (**Fig. 3P**) as compared to control cells. Together, these findings support a model in which MRGPRX4 activation promotes melanoma phenotype switching toward an ECM-remodeling, invasive MES-like state, and that this program is at least partly MRGPRX4-dependent.

Together, these spatial and transcriptional analyses reveal that MRGPRX4-driven melanoma selectively expands a mesenchymal and neural-crest-aligned invasive state that is preferentially positioned at vascular interfaces. The convergence of inflammatory signaling and loss of melanocytic identity within this compartment suggests that MRGPRX4 acts as a molecular node, possibly linking tumor-vascular interactions to invasive cell-state transitions. These findings suggest MRGPRX4 may be a determinant of melanoma progression and microenvironmental adaptation.

### MRGPRX4 promotes an invasive phenotype dominated by an aberrant PI3K-AKT pathway

We wanted to begin by examining the role of MRGPRX4 in defining melanoma cell behavior. A2058 cells that received short hairpins (sh) targeting *MRGPRX4* displayed a growth defect compared to cells receiving control sh (shLuc) (**Supplementary Fig. 4A** and **Fig. 4A**). Like the knock-down data, *MRGPRX4*^-/-^ cells had a significant proliferative defect in vitro compared to *MRGPRX4*^+/+^ cells (**Fig. 4B**). Next, to evaluate the role of MRGPRX4 *in vivo*, we utilized an immunodeficient NOD scid gamma (NSG) mouse strain that permits engraftment of human tumor cells. Consistent with *in vitro* data, *MRGPRX4*^-/-^ A2058 cells formed significantly smaller primary tumors upon implantation into the flank of immune-deficient NSG mice (**Fig. 4C**). We next assessed invasion and metastasis using both *in vitro* and *in vivo* assays. In a standardized 3D spheroid invasion assay^53^, *MRGPRX4*^-/-^ cells showed a profound loss of invasive capacity relative to *MRGPRX4*^+/+^ cells (**Fig. 4D, E**). To evaluate metastatic potential *in vivo*, we introduced GFP-labeled *MRGPRX4*^+/+^ and *MRGPRX4*^-/-^ cells intravenously into NSG mice and quantified metastatic seeding. In line with our *in vitro* findings, 6-8 weeks after injection, we found significantly less *MRGPRX4*^-/-^ cells in the lung (**Fig. 4F, G**) and liver **(Fig. 4H-J**) than *MRGPRX4*^+/+^ cells. These data suggest that MRGPRX4 can contributes to metastatic cascade, including survival, extravasation, and/or colonization of melanoma cancer cells. Overall these *in vivo* and *in vitro* data demonstrates that *MRGPRX4* can drive both proliferation and invasion in melanoma. Importantly, these differences emerged in the absence of any added exogenous ligand.

**Figure 4.**
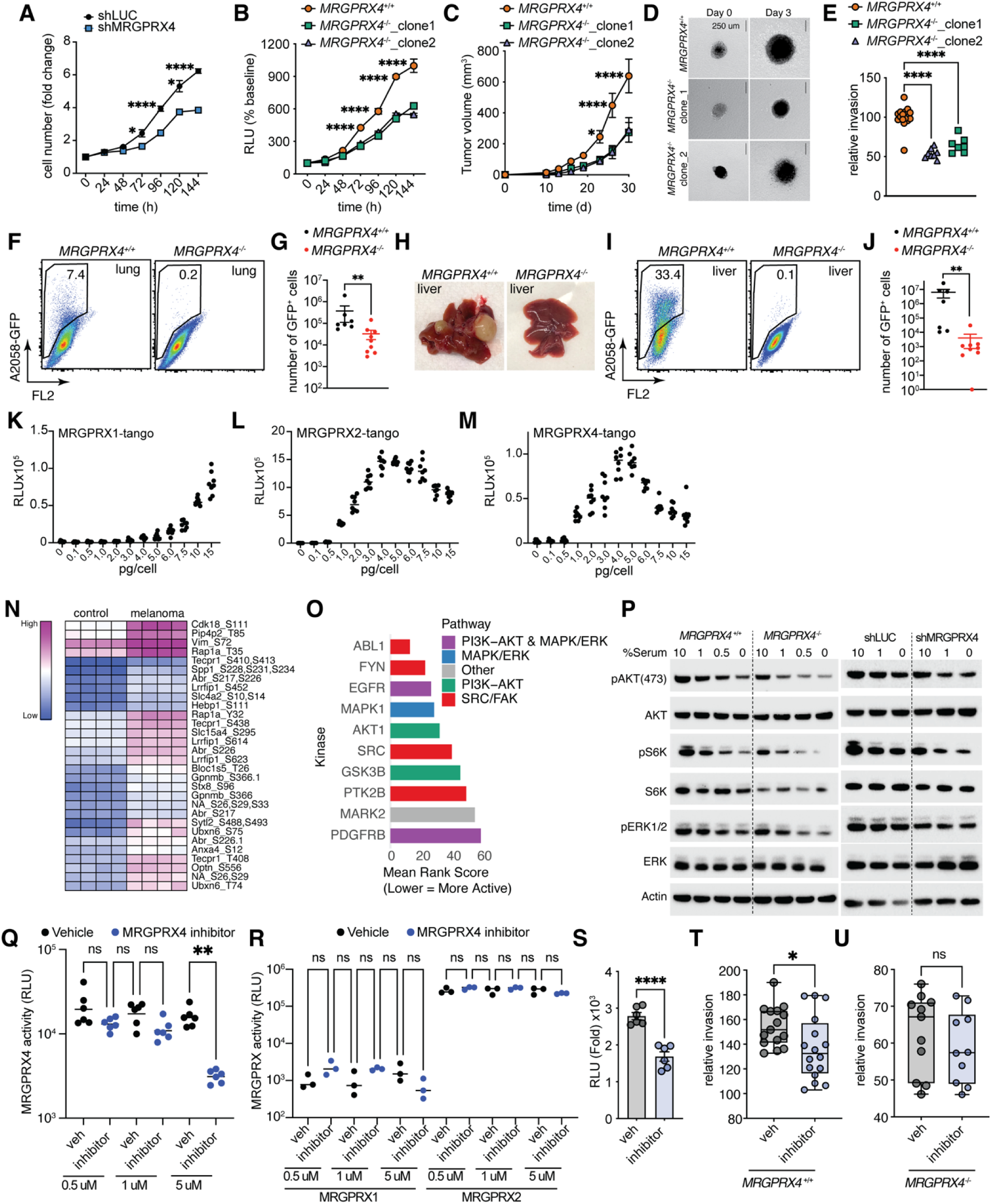
MRGPRX4 promotes an invasive phenotype dominated by an aberrant PI3K-AKT pathway. (A) Cell numbers in control and MRGPRX4 shRNA knockdown in A2058 cells. Data are mean+SEM and representative of 2 independent experiments. (B) MRGPRX4^+/+^ and MRGPRX4^-/-^ cells were cultured and ATP measurements were taken every 24 hours. Data are mean+SEM and representative of 2-3 independent experiments. (C) Tumor volumes in NSG mice (n=5-7) injected with *MRGPRX4*^+/+^ and *MRGPRX4*^-/-^ cells. Data are mean+SEM and representative of 2 independent experiments. (D-E) In vitro invasion assay. (D) Representative images (scale bar 250 um) of spheroids on days 0, 3 and (E) invasion of *MRGPRX4*^+/+^ and *MRGPRX4*^-/-^ spheroids relative to wildtype on day 3. Data are mean+SEM and pooled from 2 independent experiments. (F-J) *In vivo* invasion assay in NSG mice injected i.v. with GFP^+^ *MRGPRX4*^+/+^ or GFP^+^*MRGPRX4*^-/-^ cells and assessed 6-8 weeks later. (F) Representative flow plots, and (G) numbers of GFP^+^ cells in the lungs. (H) Representative images of the liver, (I) representative flow plots, and (G) numbers of GFP^+^ cells in the liver. Data are mean+SEM and pooled from 2 independent experiments. (K-M) Basal activity measured using PRESTO-Tango assay in HTLA cells transiently transfected with increasing doses of (K) MRGPRX1, (L) MRGPRX2 and (M) MRGPRX4. Data are mean+SEM and representative of 2-3 independent experiments. (N-O) Phosphosite enrichment in MRGPRX4-driven melanoma. (N) Heatmap of differentially regulated phosphosites in control (Tyr^CreER-^;MRGPRX4^LSL+/-^) or melanoma skin samples (Tyr^CreER+^MRGPRX4^LSL+/-)^ or (O) Kinase enrichment analysis (KEA3) in tumor samples based on phosphosite signatures. (N) heatmap nomenclature for control vs melanoma is different (P) Western blot for pAKT, pS6K and pERK1/2 in *MRGPRX4*^+/+^ and *MRGPRX4*^-/-^ A2058 cells or A2058 cells subjected to control (shLuc) or shRNA-mediated *MRGPRX4* knockdown, cultured in decreasing serum conditions. Data are representative of 2-3 independent experiments. (Q,R) Cultured HTLA cells were transiently transfected with indicated MRGPRX-Tango constructs and treated with vehicle or MRGPRX4 antagonist. (Q) MRGPRX4 and (R) MRGPRX1 and MRGPRX2 basal activity was assessed. Data are mean+SEM and representative of 2 independent experiments. (S) A2058 cells were treated with vehicle or MRGPRX4 antagonist and cellular proliferation using ATP assay was measured on Day 6. Data are mean+SEM and representative of 2 independent experiments. (T,U) A2058 cells were treated with vehicle or MRGPRX4 antagonist and invasion using the spheroid assay was assessed 48h later for (T) *MRGPRX4*^+/+^ A2058 cells and (U) *MRGPRX4*^-/-^ A2058 cells. Data are mean+SEM and pooled from 2 independent experiments. For all experiments both male and female mice were used and data are analyzed by Student’s t test or one-way ANOVA followed by post hoc test. *p < 0.05, **p < 0.01,

Because constitutive (ligand-independent) signaling is a common property of many GPCRs and can drive disease phenotypes^54-59^, we hypothesized that MRGPRX4 may promote melanoma behavior through constitutive signaling rather than classical ligand-dependent activation. To directly assess basal activity, we used the PRESTO-Tango β-arrestin recruitment assay, a widely adopted platform for quantifying GPCR signaling^60^. To measure constitutive signaling, we transfected HTLA cells with Tango constructs for MRGPRX4 and other MRGPRX family members across a range of DNA doses and quantified β-arrestin-dependent reporter activity in the absence of exogenous ligand (**Fig. 4K-M**). This revealed distinct basal signaling profiles, reflecting differential constitutive activity. MRGPRX1 exhibits little basal signaling at low expression levels (**Fig. 4K**). Still, it shows a sharp increase at high plasmid doses, indicating a receptor that may remain largely silent under physiological conditions but can activate strongly when overexpressed. This suggests that it likely requires ligand to signal efficiently. MRGPRX2 showed robust constitutive signaling across all expression levels (**Fig. 4L**), in line with work that shows an inverse agonist can inhibit basal recruitment of Gq by MRGPRX2 signaling^55^. MRGPRX4, meanwhile, exhibits a bell-shaped activity curve, with basal activity increasing at moderate expression levels but declining at higher doses (**Fig. 4M**), likely due to receptor desensitization, internalization, or modulation of surface expression via RAMP proteins^54^.

We did not want to exclude the possibility that bile acids, the currently known endogenous MRGPRX4 ligands^13,15^, could drive aberrant activity of this receptor in melanoma. To determine whether bile acids contribute to MRGPRX4 activation in melanoma, we quantified >70 bile-acid species in control versus melanoma-positive skin and in draining lymph nodes with confirmed tumor infiltration. Quantitative LC–MS analysis revealed two key findings: (i) there was no bile-acid enrichment in melanoma compared to control tissues, and (ii) bile acids were present at very low levels, typically ranging from 10 to 10 ³ nmol/mg tissue (**Supplementary Fig. 4B, C**). Deoxycholic acid (DCA), the most potent bile-acid agonist of MRGPRX4, has reported EC values in the 5–19 μM range (≈5×10 ³–1.9×10 ² nmol/mg tissue)^13,15,54^. In our dataset, the highest DCA concentration was ∼1.6×10 nmol/mg (∼0.16 μM), corresponding to a ∼30–120-fold deficit relative to the EC range (**Supplementary Fig. 4B, C**). All other bile-acid species were present at even lower levels, many of them 10^2^-to-10 -fold below these thresholds. These findings indicate that endogenous bile acids in the melanoma microenvironment are far below the concentrations required to activate MRGPRX4, even for its most potent known bile-acid ligand and are therefore unlikely to serve as physiologic drivers of MRGPRX4 signaling in our model. These findings strongly suggest that dysregulated MRGPRX4 signaling in melanoma arises from intrinsic basal activity rather than ligand-dependent activation.

To define signaling programs downstream of aberrant MRGPRX4 signaling *in vivo*, we performed unbiased phosphoproteomic profiling on dorsal skin from Tyr^CreER+^;MRGPRX4^LSL^ mice with established melanoma compared to Tyr^CreER-^;MRGPRX4^LSL^ controls. Global phosphopeptide intensity distributions were comparable between groups, indicating similar depth and sample quality (**Supplementary Fig. 4D**). In contrast, principal component analysis revealed clear segregation of tumor versus control samples (**Supplementary Fig. 4E**), consistent with broad remodeling of phosphorylation networks. As expected, most sites identified were phospho-serine, with smaller contributions from phospho-threonine and phospho-tyrosine residues (**Supplementary Fig. 4F**). Differential analysis identified a robust set of regulated phosphosites that segregated samples into distinct clusters (top 100 shown in **Supplementary Fig. 4G**). Gene ontology analysis revealed strong enrichment for actin-binding and cytoskeletal pathways, including Z-disc and sarcomere organization, cell–cell junction components, and intermediate filament assembly (**Supplementary Fig. 4H**). Functionally, these phosphorylation changes point to increased cytoskeletal activity within tumor cells, consistent with greater contractility and increased interaction with surrounding matrix. These features align with a more mesenchymal-like state and may support enhanced invasive behavior. Among the regulated phosphosites, we observed increases in proteins linked to proliferation (CDK18), cytoskeletal remodeling (Vimentin), trafficking (PIP4P2, RAP1A), and stress-responsive Rho-GTPase pathways (TECPR1, ABR, LRRFIP1) (**Fig. 4N**), reflecting a shift away from a canonical melanocytic profile toward an activated, invasive tumor state.

Kinase enrichment analysis (KEA) further revealed preferential activation of ABL1, FYN, EGFR, MAPK1, and AKT1, indicating convergence on MAPK/ERK- and PI3K-Akt-driven signaling axes (**Fig. 4O**). Notably, strong enrichment of Src-family kinases (FYN, SRC, PTK2B), together with ABL1 and PDGFRB, reflects signaling properties associated with migration, ECM engagement, and mesenchymal-like cell states that are thought to correlate with metastatic dissemination and attenuated MAPK inhibitor response^61-63^. In line with the role for PI3K-AKT-mTOR signaling in melanoma, pharmacologic inhibition of these pathways using torin or dactolisib fully abrogated A2058 proliferation (**Supplementary Fig. 4I, J**). Consistent with the *in vivo* kinase signatures, loss of MRGPRX4 signaling in A2058 melanoma cells, via gene knockout and knockdown strategies, reduced phosphorylation of key MAPK- and mTOR-dependent effectors, including pAkt, pS6K, and pERK1/2 (**Fig. 4P**), validating the direct reliance on this signaling axis. Together, these findings demonstrate that MRGPRX4 signaling drives a cytoskeletal-active, kinase-reprogrammed melanoma state by engaging aberrant PI3K-AKT/MAPK signaling.

Lastly, we asked whether MRGPRX4 could be pharmacologically targeted as a proof-of-principle approach. We used compound 31-2, a commercially available MRGPRX4 antagonist, to test whether acute inhibition of basal MRGPRX4 signaling could affect melanoma cell behavior. In PRESTO-Tango assays, compound 31-2 selectively reduced basal reporter activity in MRGPRX4-expressing cells, with the clearest effect observed at 5 μM, while having no measurable effect on MRGPRX1- or MRGPRX2-transfected cells under the same conditions (**Fig. 4Q–R**). Because this reduction occurred in the absence of exogenous agonist, these data are consistent with inverse-agonist-like suppression of constitutive MRGPRX4 signaling rather than simple competitive antagonism. These findings further support the conclusion that MRGPRX4 behaves as a basally active GPCR in melanoma cells. Because this ligand-independent activity was readily detectable, we hypothesized that melanoma cells may leverage constitutive MRGPRX4 signaling to support tumor growth and invasion. Functionally, compound 31-2 reduced proliferation and partially reduced invasion of parental A2058 melanoma cells (**Fig. 4S–T**), whereas the same treatment had no detectable effect on MRGPRX4-deficient A2058 cells (**Fig. 4U**). Although the magnitude of pharmacologic inhibition was smaller than that observed with genetic MRGPRX4 loss, this is consistent with acute and potentially incomplete receptor inhibition by a micromolar-level, non-optimized tool compound. Together, these data provide proof-of-principle that MRGPRX4 can be pharmacologically targeted and suggest that suppression of basal MRGPRX4 signaling can partially phenocopy the anti-proliferative and anti-invasive effects of genetic MRGPRX4 loss.

In summary, these data support a model in which MRGPRX4 acts as a basally active GPCR that promotes melanoma reprogramming toward a mesenchymal, invasion-competent state and engages kinase signaling networks required for tumor expansion and metastatic spread.

## Discussion

Melanoma remains the most lethal skin cancer, with a steadily rising incidence worldwide^64^. Here, we identify the itch-linked GPCR, MRGPRX4, previously considered a sensory neuron-restricted receptor, as an unexpected melanoma oncogene-like driver. We show that MRGPRX4 is selectively expressed in neural-crest-like, pre-EMT melanoma states, a transcriptional program associated with dedifferentiation, invasiveness, and therapeutic resistance. Functionally, MRGPRX4 drives proliferation, survival, and invasion *in vitro* and accelerates tumor growth and metastatic behavior *in vivo*. These findings uncover a previously unrecognized GPCR-driven axis of melanoma progression and position MRGPRX4 as a receptor with oncogenic-like properties, co-opted during tumor dedifferentiation.

Although nearly 50% of melanomas harbor BRAF mutations and initially respond to combined BRAF/MEK inhibition^65^, durable responses remain rare due to the rapid emergence of adaptive resistance^66-68^. Our work identifies MRGPRX4 as a parallel activator of MAPK and PI3K-AKT-mTOR signaling, two pathways recurrently engaged by drug-persistent melanoma cells. Prior studies show that PI3K pathway activation enables survival of MAPK-inhibited persisters^69-72^. Our data further support this interpretation. Proteomics analysis of MRGPRX4-driven melanoma revealed enrichment of insulin/IGF-1 receptor pathways, known to sustain melanoma proliferation, survival, and treatment resistance via PI3K-AKT and MAPK signaling^73,74^. Additionally, phosphoproteomic kinase-substrate mapping further implicated EGFR-associated receptor tyrosine kinase (RTK) nodes consistent with growth factor-driven MAPK/PI3K signaling and invasive properties in melanoma^75^. Notably, G-protein-coupled receptors can transactivate EGFR, providing a mechanistic framework linking aberrant GPCR signaling to RTK-dependent pathways^76^ and provides a mechanistic bridge linking a non-canonical GPCR to classical RTK-dependent signaling cascades. We therefore propose that constitutively active MRGPRX4 establishes an PI3K/RTK-connected signaling axis that reinforces proliferative and invasive phenotypes.

Importantly, melanoma arises in our *Tyr*^CreER+^;*MRGPRX4*^LSL^ model without engineered tumor-suppressor loss, suggesting that sustained MRGPRX4 signaling alone can reinforce proliferative and survival circuits, including RTK-linked cell-cycle deregulation, thus lowering the threshold for malignant initiation and progression. The robust pulmonary and hepatic colonization observed in NSG mice further suggests that MRGPRX4 confers metastatic competence in vivo, consistent with a model in which constitutive GPCR activity sustains both proliferative and invasive programs.

Our data indicate that ligand-dependent activation is unlikely to be the dominant mode of MRGPRX4 signaling in cutaneous melanoma. Although bile acids are *bona fide* MRGPRX4 agonists and can promote tumorigenesis in hepatobiliary settings^77^, quantitative lipidomics revealed no bile-acid enrichment in melanoma versus control skin, and absolute concentrations fell well below EC50 values for receptor activation. Thus, bile-acid-MRGPRX4 signaling is unlikely to contribute to primary skin manifestations. However, metastatic spread to the liver or gastrointestinal sites, where bile acids are abundant, may create a distinct context in which ligand-dependent MRGPRX4 activation could further amplify tumor growth, consistent with the pronounced hepatic tumor burden observed *in vivo*.

MRGPRX4 also selectively reshapes the tumor microenvironment. MRGPRX4-driven tumors contained fewer eosinophils but increased neutrophils, together with an expanded pool of PD-L1 macrophages and neutrophils, and elevated PD-L1 on CD45 tumor-associated cells. In contrast, PD-L1 remained unchanged on dendritic cells and monocytes. This represents a selective immune shift characterized by a PD-L1-rich myeloid and tumor-stromal niche that could be predicted to suppress T cell responses and enhance tumor persistence.

Finally, our findings connect melanoma dedifferentiation to developmental ontogeny. Because melanocytes and sensory neurons both arise from neural crest lineages, re-expression of MRGPRX4 in neural-crest-like melanoma states suggests that dormant lineage programs become aberrantly reactivated. Environmental modulators, such as UV exposure, inflammatory cytokines, or an aging-associated extracellular milieu, may converge on transcriptional nodes permissive for MRGPRX4 expression. In this model, melanoma co-opts a neuron-restricted GPCR to acquire proliferative and invasive properties, suggesting that MRGPRX4 blockade is a rational lineage-restricted therapeutic strategy.

## Limitations

While our mouse model shows comparable tumor formation in both sexes, epidemiologic data indicate sex-linked differences in melanoma immunity and progression^78^. Future studies should determine whether MRGPRX4 expression or signaling varies across age or sex in patients.

Additionally, while MRGPRX4 shows enhanced expression in patients, the precise epigenetic mechanisms that govern this biology remain to be elucidated.

## Acknowledgments

This work is supported by the Howard Hughes Medical Institute support to X.D. X.D. is also supported by R37NS054791. S.L. is supported by NIH grants R01AI170709. J.P. is supported by American Lebanese Syrian Associated Charities (ALSAC) of St. Jude Children’s Research Hospital. Y.C. is supported by Pardee Foundation and Fox Chase Cancer Center. We thank Ted Choi at 3D Genomics for helping with melanoma spatial transcriptomics studies. We thank Hao Zhang and the Bloomberg Flow Cytometry and Immunology core for cell sorting. We thank the Department of Molecular Microbiology and Immunology Microscopy Facility at the Johns Hopkins Bloomberg School of Public Health for support and use of their instruments, including the Leica STELLARIS 8 FALCON confocal microscope (supported by NIH S10OD036404). We thank the Johns Hopkins Genetic Resources Core facility for help with bulk RNA sequencing.

## Author contributions

N.G. performed experiments with assistance from A.A., H.M.Y., A.M., and H.J. A.A. provided confocal imaging data. M.A, M.L. provided data with shRNA knockdowns (proliferation) and proteomics (Western blots). Z.W., and J.P. provided skin total proteomics and phospho-proteomics data. Y.C. and A.W. helped with melanoma assays and melanoma expertise. S.L. performed bioinformatics analysis (RNA-seq, scRNA-seq, proteomics, phosphoproteomics, spatial transcriptomics, epigenetics). N.G., S.L. and X.D. conceptualized the study and wrote the paper.

## Methods

### Mice

MRGPRX1^R26-stop-flox^ (MRGPRX1^LSL^), MRGPRX2^R26-stop-flox^ (MRGPRX2^LSL^), MRGPRX4^R26-stop-flox^ (MRGPRX4^LSL^) animals were generated as previously described^13,18,19^. The MRGPRX3^R26-stop-flox^ line was generated by homologous recombination of an MRGPRX3 cDNA construct under loxp-STOP-loxp control to the ROSA26 locus. NSG mice (stock#005557) and Tyr^CreER^ animals were obtained from Jax (stock# 012328). *Tyr*^CreER^ animals were crossed to MRGPRX^LSL^ animals and at weaning (3-4 weeks), the experimental and control animals were given 1 mg/40g i.p.^30^ injections of tamoxifen in corn oil for 3 consecutive days to induce melanoma. For experiments, both male and female mice were used. All mice were maintained in a specific-pathogen free facility and used according to the Institutional Animal Care and use Committee.

### Real-time PCR

Total RNA from skin tumor samples or A2058 cells was harvested using TRIzol (Thermo) and used for synthesis of cDNA (Applied Biosystems). PCR was performed using primer probes (Thermo). As *MRGPRX4* is a single exon gene, total RNA was treated with ezDNAse (Thermo) and cDNA generated with or without reverse transcriptase (Thermo). Gene expression for *MRGPRX4 (Hs00607779)* was done using TaqMan probes (Thermo) and data are expressed as relative expression to the housekeeping gene ribosomal protein S13 (*RPS13*) or *GAPDH*.

### Generation of *MRGPRX4* ^-/-^ A2058 cells

CRISPR-Cas9-mediated knockout cell clones of MRGPRX4 were generated commercially (EditCo Bio, Inc.) using a pooled transfection strategy such that multiple edited alleles were initially represented. A non-targeting control-transfected cell pool was generated in parallel (EditCo Bio, Inc.) and used throughout experiments. Following editing, single-cell isolation was performed using a single-cell dispensing system (EditCo Bio) and deposited into 96- or 384-well plates at 1 cell/well. Plates were imaged every 3 days to confirm clonal expansion from a single founder cell. For each putative clone, genomic DNA was isolated and the edited MRGPRX4 locus was PCR-amplified and subjected to Sanger sequencing. Editing outcomes at the sgRNA target region were analyzed using the ICE deconvolution pipeline (EditCo Bio), revealing frameshift-inducing indels predicted to generate premature stop codons in *MRGPRX4*. Two independent knockout clones (clone_1 and clone_2) with distinct indel patterns were selected for downstream characterization. The sgRNA used for targeting human *MRGPRX4* had the protospacer sequence 5′-AACGTATAATCTGGAAGCTG-3′ with a canonical TGG PAM. This target lies within the single coding exon of *MRGPRX4*.

### *In vitro* proliferation assay

A2058 cells were seeded at a density of 500-1000 cells in 100 uL in complete DMEM (10% FBS) in black, flat-bottom 96-well plates. Proliferation was assayed using Promega CellTiter-Glo2.0 ATP assay according to the manufacturer’s protocol. Briefly, cells were equilibrated at RT for 30 minutes before the addition of luminescent reagent for 10 minutes and signal quantified using a luminescent microplate reader. Data are calculated as a relative percentage of the ATP reading from day 0. For experiments utilizing the MRGPRX4 antagonist, cells were treated with vehicle (DMSO) or 5 uM of the MRGPRX4 antagonist (HY-145997, MedChemExpress) twice every day. For experiments using chemical inhibitors of the PI3K-mTOR pathways, torin 1 (HY-13003, MedChemExpress) and dactolisib (HY-50673, MedChemExpress), cells were treated with vehicle or inhibitors (50 nM) on day 0, and proliferation was determined on the following days.

### NSG-xenograft model

A total of 0.5X10^6^ A2058 *MRGPRX4* ^+/+^ and *MRGPRX4* ^-/-^ cells were injected subcutaneously in the flank of NSG mice. Tumor growth was measured using Mitutoyo digital calipers and two measurements (length and width) per growth were taken. Tumor volume was calculated using the formula (0.5 x L x W^2^). Mice with tumors >1.5 cm in any dimension were euthanized.

### *In vivo* metastasis model

A total of 2.5 × 10 A2058 *MRGPRX4* ^+/+^ and *MRGPRX4* ^-/-^ cells were seeded per T182 cm² tissue culture flasks the evening prior to transduction. The following morning (Day 0), cells were transduced with a GFP-expressing lentivirus (VectorBuilder) at a multiplicity of infection (MOI) of 1 in the presence of 8 μg/mL polybrene. Twenty-four hours post-transduction (day 1), the media was removed and replaced with fresh complete media (DMEM+10% FBS). On day 3 (72 hours post-transduction), cells were harvested via trypsinization and GFP cells were flow-sorted (DAPI^-^GFP^+^). 2.5×10 GFP cells were resuspended in 200 μL PBS and injected intravenously via tail vein into NSG mice. 6-8 weeks following injection, the lungs and livers were harvested and enzymatically dissociated using the Miltenyi Tumor Dissociation Kit, according to the manufacturer’s protocol. GFP cells were then quantified by flow cytometry.

### Flow cytometry

Control, tumor skin, lung and liver tissue were dissociated using the Miltenyi Tumor Dissociation Kit according to the manufacturer’s protocol. Following tissue digestion, cells were processed for RBC lysis using ACK lysis buffer. Recovered cells were counted, plated at a density of 1-2 X 10^6^ cells in a 96 well round-bottom plate and processed for flow staining as previously described^79^. Briefly, cells were stained with the live/dead Zombie-Aqua stain for 10 min (RT) and blocked with 20 ug/ml anti-CD16/32 (BioXCell) for an additional 20 min (RT). Cells were stained with fluorochrome-labeled antibodies (CD45-clone 30-F11, CD64-clone S18017D, MerTK-clone 2B10C42, MHCII-clone M5/114.15.2, CD11c-clone N418, Ly6G-clone 1A8, Siglec F- clone E50- 2440, CD11b-clone M1/70, Ly6C clone- HK1.4, CD317-clone eBio927, PD-L1-clone M1H7). Data were acquired on an LSRII flow cytometer (BD Biosciences) and gated to exclude debris and select single cells (SSC-W/SSC-A). Data were analyzed using FlowJo (BD Biosciences).

### TCGA/GTEx/cBioPortal expression data

Gene expression data from normal skin and melanoma samples were analyzed using the UCSC Xena Browser (https://xena.ucsc.edu). Bulk RNA-Seq data for normal skin tissue were obtained from the Genotype-Tissue Expression (GTEx) and melanoma tumor expression data were sourced from The Cancer Genome Atlas (TCGA). For each tumor type, we calculated the median expression and used the Wilcoxon rank-sum test to compare the median values with those of all other samples. Median differences were plotted against minus log10 (p-values). Bulk RNA-seq data for normal human skin melanocytes^80,81^, keratinocytes^81^ and fibroblasts^81^ were downloaded from cBioPortal. For cross-species projection of the mouse proteomics-derived signature onto human melanoma tumors, TCGA-SKCM gene expression quantification files were downloaded from the NCI Genomic Data Commons.

### Immunofluorescence of mouse tissue

Mouse melanoma tissue samples were embedded in Tissue-Tek O.C.T Compound (Sakura Finetek, 4583), snap frozen and stored in -80°C until cutting. Sections of 10um were cut using a Microm HM560 Cryostat and were treated using HistoVT One (Nacalai Tesque, Kyoto, Japan) at 70°C for 30 min, followed by cooling at RT before immunostaining, as previously described^82^. Donkey serum was used to prepare blocking buffer and antibody cocktails. Sections were washed with 0.5% Triton-X100 prepared in 1X PBS and then blocked for 1-2 hours at RT. After blocking, the primary antibodies: MRGPRX4 (Invitrogen, PA5-33954, 1:100), PRRX1 (NovusBio, NPB1-06067, 1:200), CD31 (BD Biosciences, 550274, 1:00) and PROX1 (NovusBio, NBP1-30045, 1:100) were incubated overnight at 4°C. Before adding secondary antibodies, the sections were washed three times with 0.5% Triton-X100/PBS. The secondary antibodies: donkey anti-goat Alexa Fluor 647 (Invitrogen- A21447), donkey anti-rat Alexa Fluor 594 (Invitrogen- A21209), donkey anti-rabbit Alexa Fluor 546 (Invitrogen- A10040), donkey anti-mouse Alexa Fluor 488 (Invitrogen- A-21202), and Hoechst 33342 were incubated for 1-2 hours at room temperature, followed by 0.5% Triton-X100/PBS washes. Samples were mounted using ProLong™ Glass Antifade Mountant (Thermo Fisher Scientific, Waltham, MA, USA) and allowed to cure overnight before imaging.

### Immunofluorescence of paraffin-embedded patient skin tissue

Formalin-fixed, paraffin-embedded (FFPE) melanoma patient skin samples were obtained commercially. Patient characteristics are as follows, 1 male (68yr., Caucasian, stage IIIC) and 1 female (64yr. Caucasian, stage II). Sections (5 um) of FFPE were baked at 60°C for 2 hours before proceeding to deparaffinization. Briefly, FFPE sections were dewaxed in fresh xylene, followed by graded ethanol washes and rinsed in nuclease-free water. Antigen retrieval was performed using HistoVT One (Nacalai Tesque, Kyoto, Japan) at 70°C for 30 min, followed by cooling at RT before immunostaining. Donkey serum was used to prepare blocking buffer and antibody cocktails. Sections were washed with 0.5% Triton-X100 prepared in 1X PBS and then blocked for 1-2 hours at RT. After blocking, anti-MRGPRX4 (Invitrogen, PA5-33954, 1:100), anti-SOX-10 (Invitrogen, PA5-37890, 1:100), and anti-HMB-45 (recognizes gp100/PMEL, Invitrogen, MA1-34759, 1:100) were incubated overnight at 4°C. Before adding secondary antibodies, the sections were washed three times with 0.5% Triton-X100/PBS. The secondary antibodies: donkey anti-goat Alexa Fluor 647 (Invitrogen- A21447), donkey anti-rabbit Alexa Fluor 546 (Invitrogen- A10040), donkey anti-mouse Alexa Fluor 488 (Invitrogen- A-21202), and Hoechst 33342 were incubated for 1-2 hours at room temperature, followed by 0.5% Triton- X100/PBS washes. Samples were mounted using ProLong Glass Antifade Mountant (Thermo Fisher Scientific, Waltham, MA, USA) and allowed to cure overnight before imaging.

### Fluorescence Imaging and Processing

Images were acquired using a Leica Stellaris 8 Falcon confocal microscope (Leica Microsystems, Germany). All images were acquired at 1024 × 1024 pixels and 16-bit depth. Detector gain, laser power, and pinhole settings were adjusted to minimize photobleaching and maximize signal-to-noise ratio. Post-acquisition visualization and linear contrast adjustments were performed in LAS X software (Leica). Confocal TIFF images were batch-processed using a custom Python pipeline (Python 3.12.12; scikit-image, SciPy, NumPy, pandas, matplotlib). Nuclei masks were generated using standard image segmentation, and marker intensities (MRGPRX4, PRRX1) were quantified per nucleus; cells were classified as positive or negative for each marker using adaptive per-image intensity thresholds. Blood/lymphatic vessel structures were identified from CD31 and PROX1 channels using vessel-enhancement filtering and morphological cleanup, and vessel-adjacent regions were defined by dilating the vessel mask by 5 µm. Cells were annotated as vessel-associated (BV/LV^+^) or non-vessel-associated (BV/LV^-^) based on centroid proximity to the vessel mask. Per-nucleus measurements and per-region percentages were exported as CSV files. Paired comparisons between BV/LV^+^ and BV/LV^-^ regions were performed using Wilcoxon signed-rank test (two-tailed), and data were visualized using box-and-scatter plots.

### 3D Spheroid Assays

Melanoma cells (5000 cells/well) were plated in 1.5% agar, and spheroids were allowed to form for 3 days in a 96-well plate followed by embedding in rat-tail collagen I (Thermo Scientific) as previously described^53^. For experiments using the MRGPRX4 antagonist, cells were treated with vehicle (DMSO) or the antagonist (10 uM) twice daily. All spheroids were imaged at time point 0 (t0) and later (t1) time points post-embedding. The spheroids were imaged using a Nikon Eclipse Ti2 inverted microscope. Quantitation of invasive surface area was performed in a blinded fashion using Image J and percent invasion for a spheroid is calculated relative to day 0 using the formula ((t1-t0)/t0) *100.

### Spatial transcriptomics

#### Tissue processing

Lower back skin was depilated, and tumor tissue was excised. Tissues were embedded in OCT and frozen in an isopentane/liquid nitrogen bath. Blocks were kept at -80°C until processing.

#### Spatial transcriptomics data generation and preprocessing

Serial 10 µm frozen sections of mouse melanoma were mounted on Visium Spatial Gene Expression slides (10x Genomics) and stained with H&E. Libraries were prepared using the Visium Spatial protocol and sequenced on an Illumina NovaSeq 6000. Raw FASTQ files were processed with Space Ranger v1.3.1 to align reads to the mm10 genome and generate spot level UMI count matrices. The resulting data were imported into R (v4.2.2) via Seurat v4.3.0, and low quality spots (fewer than 200 detected genes or >10% mitochondrial reads) were excluded. Data were log-normalized and the top 2,000 variable genes identified for downstream analyses.

#### Bayesian sub-spot resolution enhancement

To improve the 55 µm diameter limitation of individual Visium spots, we used BayesSpace v1.6.0 for Bayesian spatial resolution enhancement. Data was first preprocessed (spatialPreprocess()) using log normalized counts and principal component analysis on the top 15 PCs. A spatial neighbor graph was constructed with default parameters. Spot level clustering into seven spatial domains (q = 7) was performed by Markov Chain Monte Carlo (spatialCluster(), 30,000 iterations, γ = 3). We then subdivided each spot into a 3 × 3 grid of pseudospots (∼18 µm resolution) via spatialEnhance() (10,000 total iterations with 2,000 burn in, γ = 3). Enhanced clusters and expression matrices were stored in a SingleCellExperiment object for subsequent mapping.

#### Gene-set visualization

Gene signatures, like previously published^17^ were scored in each pseudospot using AUCell. For each pseudospot, raw counts were rank ordered using AUCell v1.16.0 (AUCell_buildRankings()), and area under the curve enrichment scores computed (AUCell_calcAUC()) with an aucMaxRank cutoff of 5% of genes. AUC scores were appended to pseudospot metadata and visualized in ggplot2 v3.4.2.

### Dimensionality reduction and clustering of snRNAseq data

snRNA-seq data were normalized using SCTransform and analyzed by PCA. A shared nearest neighbor graph was constructed using PCA embeddings (FindNeighbors) and clustered using the Louvain algorithm (FindClusters) at a resolution yielding 19 clusters, which were visualized using UMAP (RunUMAP). Melanoma cells were further subsetted for analysis.

### Public scRNA-seq datasets

#### Human cutaneous melanoma scRNA-seq

To assess *MRGPRX4* expression in human cutaneous melanoma, publicly available melanoma scRNA-seq data were obtained from GSE269936. Downstream analysis was restricted to three *MRGPRX4*-expressing samples with the strongest detectable *MRGPRX4* signal (GSM8330651, GSM8330662, and GSM8330659).

#### Comparative integration of MRGPRX4 tumors with NRAS- and BRAF-driven melanoma GEMMs

To compare melanoma transcriptional states in the MRGPRX4 model with established genetically engineered melanoma models, we integrated our tumor snRNA-seq dataset with publicly available single-cell melanoma data from GSE207592, including *Tyr::NRAS*^Q61K/°^;*Ink4a*^−/−^ and *Tyr*^CreER^;*Braf*^V600E^;*Pten*^fl/fl^ tumors. The final integrated melanoma object was analyzed in Seurat and contained 84,168 melanoma cells/nuclei. Dimensionality reduction was performed using Harmony, and integrated visualization was performed using the Harmony-corrected UMAP embedding. Tumor cells were grouped by genotype/model of origin, and melanoma transcriptional states were annotated using curated melanoma state programs, including melanocytic, IFN/antigen-presentation-associated, cycling, NC-like invasive, ECM-rich mesenchymal, MES/pre-EMT, inflammatory fibroblast-like, OxPhos/neural-leaning, and transitory neural crest/stress-associated states.

### Heatmap

Cluster markers were defined with FindAllMarkers. Top genes per melanoma type were displayed in a heatmap of average scaled expression (pheatmap).

### Gene signature scoring and visualization

Single-cell signature enrichment was quantified using UCell, a rank-based method for robust per-cell gene set scoring, computed on log-normalized RNA expression values. UCell scores for melanocytic, mesenchymal-like, cycling, and interferon-response signatures were calculated for each cell and stored as metadata, then visualized by projection onto the UMAP embedding used for melanoma state visualization. To improve separation between partially overlapping melanocytic and mesenchymal-like programs, consistent with the presence of hybrid or transitional melanoma states described in prior work, we additionally computed a mesenchymal specificity axis defined as the difference between mesenchymal-like and melanocytic scores (Mesenchymal_like_UCell−Melanocytic_UCell). This composite metric emphasizes relative program dominance rather than absolute enrichment and was rescaled to the [0,1] range for visualization.

### DepMap data analysis

Gene expression data was downloaded directly from DepMap and analyzed for MRGPRX4 expression. Further, human melanoma cell lines were grouped into MRGPRX4^hi^ ((log2(TPM+1)>1) and MRGPRX4^lo^ ((log2(TPM+1)>0.3). Genes were analyzed for pathway enrichment using over-representation analysis (ORA) based on curated REACTOME pathway annotations.

### Mutation analysis in DepMap and TCGA

We assessed MRGPRX4 mutation frequency across cancer using DepMap and TCGA. For DepMap, we used the public mutation dataset and the “Damaging Mutations” field to identify cell lines carrying protein-disrupting *MRGPRX4* variants and compared melanoma lines with all other lineages. For TCGA, we analyzed the publicly released MC3 pan-cancer MAF. Mutations in *MRGPRX4* were extracted by gene symbol and classified using the MC3 Variant_Classification annotation. We defined protein-altering damaging mutations as missense, nonsense, frameshift, in-frame indels, splice-site, or start/stop-codon alterations, and excluded synonymous/non-coding variants. For each TCGA cancer type, we counted unique tumors harboring at least one protein-altering *MRGPRX4* mutation and compared melanoma with all other cancers.

### MRGPRX4 promoter methylation analysis

Public Illumina HumanMethylation450K data from GSE120878 were analyzed. β-values were obtained from series matrix files, and promoter-associated CpG probes were defined based on standard Illumina 450K annotations (hg19) within ±2 kb of the *MRGPRX4* transcription start site. Probes annotated to TSS1500, TSS200, 5′UTR, or 1stExon were retained, and median β-value across probes was used to represent promoter methylation per sample. Group-wise methylation differences were visualized, and statistical significance was assessed using a two-sided Wilcoxon rank-sum test.

### Bulk RNA sequencing

RNA from *MRGPRX4* ^+/+^ and *MRGPRX4* ^-/-^ A2058 cells was extracted using the RNAeasy kit (Qiagen) with on-column genomic DNA digestion. Library construction and sequencing were performed by the Johns Hopkins Genetic Resources Core Facility. Reads were mapped to GRCh38, and differential expression was performed using DESeq2. Data are expressed as normalized transcripts per million.

### PRESTO-Tango assay

We used HTLA cells, a HEK293 cell line stably expressing a tTA-dependent luciferase reporter and a β-arrestin-TEV (Tobacco Etch Virus) protease fusion gene for the assay.^60^ HTLA cells were grown in antibiotic containing media. HTLA cells (1x10^4^) are seeded in poly-L-lysine coated 96-well tissue culture plates. The next day cells were transiently transfected with MRGPR-Tango constructs. The following day, cells were washed twice with 1X PBS, and fresh media (DMEM / F12, 0.1% FBS) is added. After 22-24hr, supernatants were discarded and the cells were lysed in passive lysis buffer (Promega), and luciferase activity is determined using the luciferase assay system (Promega) on a BioTek Synergy Microplate reader. For experiments with MRGPRX4 antagonist, cells were transfected (2pg/cell) with MRGPR-Tango constructs and treated with vehicle or antagonist twice a day.

### Bile acid quantification

Bile acids were commercially (Creative Proteomics) quantified using targeted LC–MS/MS. Tissue homogenates (skin and draining lymph node; 10 mg input) were prepared in water with an internal standard (ursodeoxycholic acid-d4), extracted in acetonitrile, clarified by centrifugation, dried under nitrogen, and reconstituted in 50% acetonitrile. An Agilent 1290 UHPLC coupled to an Agilent 6495B triple-quadrupole mass spectrometer was operated in negative-ion MRM mode. Chromatographic separation was performed on a Waters BEH C18 column (2.1×150 mm, 1.7 μm) using a binary gradient of 0.01% formic acid in water and 0.01% formic acid in acetonitrile. Quantification was achieved using 10-point internal-standard calibration curves generated from mixtures of 74 authentic bile acid standards. Analyte concentrations were calculated from analyte-to-IS peak area ratios using linear regression.

### Mass spectrometry-based proteomics and phosphoproteomics

Proteomics profiling was carried out by the optimized tandem mass tag (TMT) strategy coupled with two-dimensional liquid chromatography and tandem mass spectrometry (LC/LC-MS/MS)^83,84^. Dissected melanoma tissues from mouse skin were homogenized in lysis buffer containing 50 mM HEPES (pH 8.5), 8 M urea, 0.5% sodium deoxycholate (SDC), and phosphatase inhibitor cocktails (Roche). Homogenization was performed using a Bullet Blender with bead-beating at speed 6 for 30 s with 10 s intervals at 4 °C for six cycles. Proteins were digested by LysC and trypsin, and the resulting peptides were reduced and alkylated, desalted using C18 solid-phase extraction (SPE) columns, and dried under vacuum.

For TMTpro labeling, peptides were resuspended in 50 mM HEPES (pH 8.5) and labeled as previously described^85^. After labeling, samples were combined in equal amounts and fractionated by offline basic pH reversed-phase LC into 40 fractions using two concatenated Waters XBridge C18 HPLC columns (4.6 mm × 250 mm, 3.5 µm). A portion of each fraction was retained for whole-proteome analysis, while the remaining material was concatenated into 20 fractions for phosphopeptide enrichment.

Phosphopeptides were enriched sequentially using both TiO₂ and Fe-NTA beads as described previously^85^. For TiO_2_ enrichment, ∼200 µg of peptides was dissolved in 30 µL binding buffer (65% acetonitrile, 2% TFA, 1 mM KH₂PO₄) and incubated with TiO₂ beads at a 3:1 bead:peptide ratio (w/w) using end-over-end rotation for 20 min at 21 °C. Beads were collected by brief centrifugation, washed twice with 200 µL washing buffer (65% acetonitrile, 0.1% TFA), and transferred onto C18 StageTips (Thermo Fisher Scientific). Phosphopeptides were eluted from TiO₂ beads using 15 µL elution buffer (15% NH₄OH, 40% acetonitrile). For Fe-NTA enrichment, TiO₂ flow-through peptides were dried, resuspended in 80% acetonitrile/0.1% TFA, and incubated with 20 µL Fe-NTA beads. Phosphopeptides were eluted using 15 µL elution buffer (5% NH₄OH, 50% acetonitrile). All phosphopeptide eluates were dried under vacuum and stored at −80 °C.

For mass spectrometric analysis, peptides were resuspended in 5% formic acid and separated on a 75-µm × 20-cm column packed with 1.7-µm C18 resin (CoAnn Technology) coupled to a Q Exactive HF Orbitrap MS. Whole-proteome samples were analyzed at 65 °C using a 14–55% buffer B gradient over 75 min (buffer A: 0.1% formic acid, 3% DMSO in water; buffer B: 0.1% formic acid, 3% DMSO in 67% acetonitrile) at a flow rate of 0.25 µL/min. Phosphoproteome samples were analyzed under similar conditions except using a 10–55% buffer B gradient over ∼120 min. Data were acquired using data-dependent acquisition, consisting of one full MS1 scan followed by ∼20 MS/MS scans. MS1 scans were collected at 60,000 resolution (AGC target 1 × 10 ; maximum injection time 50 ms). MS2 scans were acquired at 60,000 resolution with a fixed first mass of 120 m/z, scan range of 460–1600 m/z, AGC target 1 × 10, maximum injection time 110 ms, and ∼15 s dynamic exclusion.

TMTpro datasets were processed using the JUMP software suite^86^. Raw files were searched against a composite target-decoy mouse protein database (59,423 entries) derived from Swiss-Prot, TrEMBL, and UCSC. Search parameters included precursor and fragment ion mass tolerances of ±15 ppm, full tryptic specificity, up to three variable modifications, and up to two missed cleavages. Static modifications included cysteine carbamidomethylation (+57.02146 Da) and TMTpro tags on lysine residues and peptide N-termini (+304.20714 Da). Dynamic modifications included methionine oxidation (+15.99491 Da) and serine/threonine/tyrosine phosphorylation (+79.96633 Da). Peptide-spectrum matches (PSMs) were filtered based on mass accuracy, precursor ion charge clustering, and JUMP-based matching scores (J-score and ΔJn) to achieve <1% false discovery rate (FDR) at the protein or phosphopeptide level. Quantification of peptides or proteins was achieved by summarizing the intensities of reporter ions from identified peptides in the MS data.

### Cross-species comparison of MRGPRX4-driven tumor proteomics with TCGA-SKCM transcriptomes

To assess whether *Tyr*^CreER+^;*MRGPRX4*^LSL^ mouse melanoma skin tumors is reflected in human melanoma, we projected a mouse proteomics-derived gene signature onto TCGA-SKCM tumor RNA-seq expression profiles. Mouse proteins that changed between groups were mapped to their human gene symbols using orthology mapping, and genes were split into two sets: those increased in mouse tumors (“UP”) and those decreased (“DOWN”). For TCGA-SKCM tumors, gene expression values were standardized by z-scoring each gene across tumors. A per-tumor mouse program score was computed as the mean z-scored expression of mouse “UP” genes minus the mean z-scored expression of mouse “DOWN” genes. Tumors were ranked by this score and stratified by quartiles; downstream comparisons focused on the highest and lowest quartiles (Q4 vs Q1). For each gene, a TCGA Δ-expression statistic was calculated as the difference in mean expression between Q4 and Q1 tumors (Q4 − Q1). To quantify cross-species concordance, we compared mouse proteomics log2 fold-changes to TCGA Δ-expression for genes present in both datasets. We considered a gene concordant if it increased (or decreased) in both mouse and SKCM. We also summarized overall agreement using Spearman correlation between mouse log2 fold-changes and TCGA Δ-expression for the shared gene

### Western blot analysis

Cells were washed twice with phosphate-buffered saline (PBS) and lysed in a Triton X-100– based buffer containing 50 mM HEPES (pH 7.4), 2 mM EDTA, 10 mM sodium pyrophosphate, 10 mM sodium glycerophosphate, 40 mM NaCl, 50 mM NaF, 2 mM sodium orthovanadate, and 1% Triton X-100 supplemented with EDTA-free protease inhibitors (1 tablet per 25 mL). Lysates were incubated at 4°C with rotation for 10 min and clarified by centrifugation for 10 min. Protein concentrations were determined using the Bradford assay. Equal amounts of protein were mixed with sample buffer, denatured at 95°C for 10 min, resolved by SDS-PAGE using precast gels (Life Technologies), and transferred onto polyvinylidene difluoride (PVDF) membranes. Membranes were blocked with 5% non-fat dry milk in PBS containing 0.1% Tween-20 and incubated overnight at 4°C with primary antibodies. Primary antibodies used in this study were phospho-AKT (Ser473; 1:1000; CST, 9271S), total AKT (1:1000; CST, 4691S), phospho-S6K (Thr389; 1:1000; CST, 9205S), total S6K (1:1000; CST, 9202S), total ERK1/2 (1:1000; CST, 9102), and phospho-ERK1/2 (Thr202/Tyr204; 1:1000; CST, 9101), and actin (1:2000; Cell Signaling Technology, 4967S). HRP-conjugated secondary antibodies were obtained from Cell Signaling Technology (1:5000; #7074 and #7076). Chemiluminescent signals were acquired using a ChemiDoc MP imaging system.

### shRNA constructs

Short hairpin RNA (shRNA) constructs were obtained from The RNAi Consortium (TRC) library at the Broad Institute. The following TRC clones were used: sh_Luciferase (TRCN0000072254), and sh_MRGPRX4 (TRCN0000011773).

### Lentiviral production and transduction

Lentiviral particles were produced using psPAX2 and pMD2.G packaging plasmids. HEK293T cells were transfected with the indicated lentiviral constructs, and virus-containing supernatants were collected 48 h after transfection and filtered through 0.45-μm membranes. Target cells were transduced for 24 h in the presence of polybrene (8 μg/mL), after which the medium was replaced. A2058 cells were selected with puromycin (5 μg/mL) or blasticidin (20 μg/mL), depending on the lentiviral construct.

### Data availability

Spatial transcriptomics (Visium) data generated in this study have been deposited in the Gene Expression Omnibus (GEO) under accession GSE313828. Single-nucleus RNA-sequencing data have been deposited in the European Nucleotide Archive (ENA) under study accession ERP186578 (BioProject PRJEB105375), with individual run accessions ERR16022242, ERR16022341, and ERR16022347. The bulk RNA-sequencing data from A2058 melanoma cells have been deposited in GEO under accession GSE314187. The mass spectrometry proteomics and phosphoproteomics data have been deposited to the ProteomeXchange Consortium via the PRIDE partner repository with the dataset identifier PXD071277.

### Materials Availability Statement

All unique reagents generated in this study are available from the corresponding author upon reasonable request and subject to institutional Material Transfer Agreement policies.

