## Supplemental Figures for "An Itch Receptor Drives Melanoma"

Affiliations

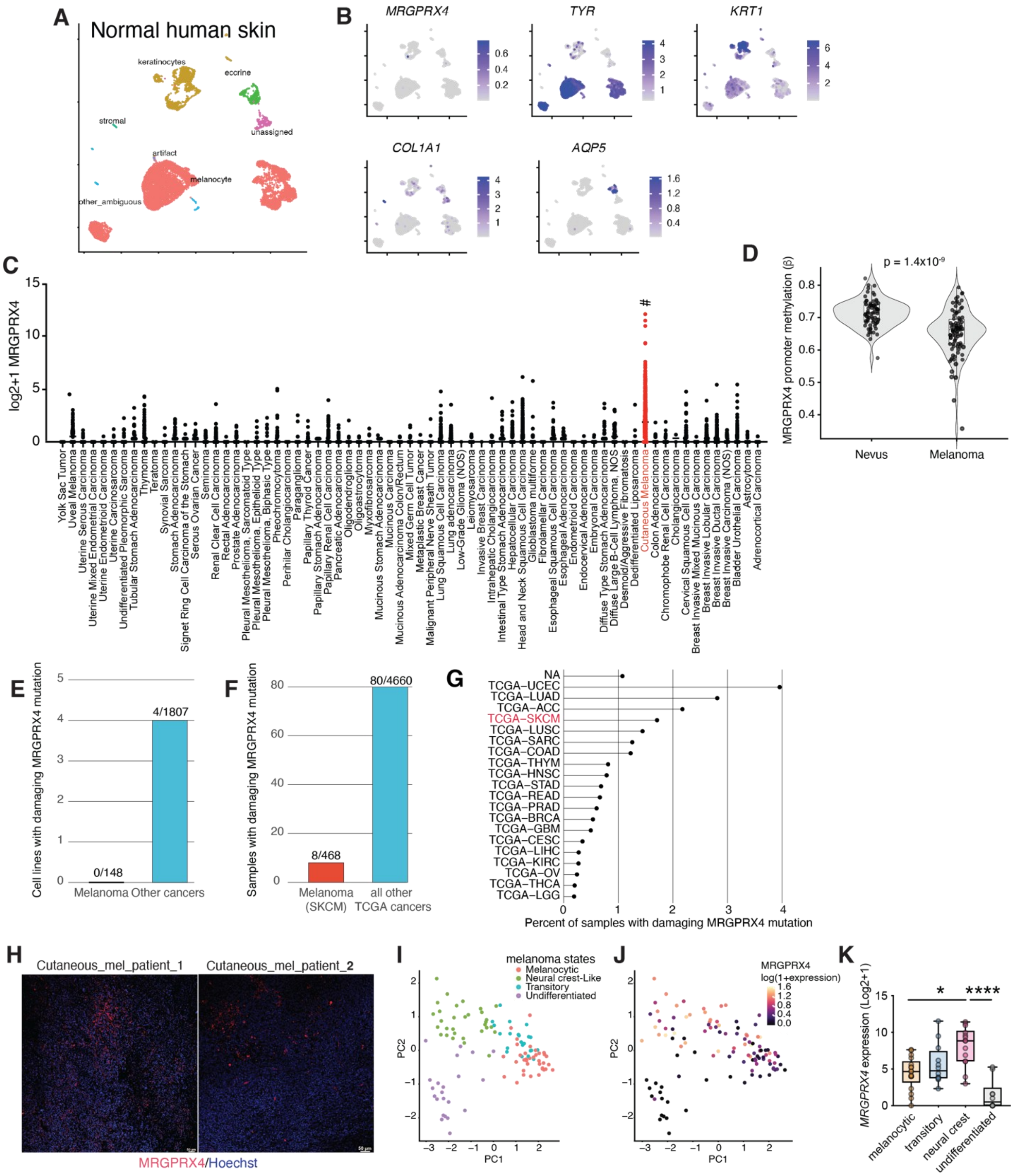
A
Normal human skin
keratinocytes
eccrine
stromal
unassigned
artifact
melanocyte
other_ambiguous
B
MRGPRX4
TYR
KRT1
COL1A1
AQP5
C
log2+1 MRGPRX4
#
Yolk Sac Tumor
Uveal Melanoma
Uterine Serous Carcinoma
Uterine Mixed Endometrial Carcinoma
Uterine Endometrioid Carcinoma
Uterine Carcinosarcoma
Undifferentiated Pleomorphic Sarcoma
Tubular Stomach Adenocarcinoma
Thymoma
Teratoma
Synovial Sarcoma
Stomach Adenocarcinoma
Signet Ring Cell Carcinoma of the Stomach
Serous Ovarian Cancer
Seminoma
Renal Clear Cell Carcinoma
Rectal Adenocarcinoma
Prostate Adenocarcinoma
Pleural Mesothelioma, Sarcomatoid Type
Pleural Mesothelioma, Epithelioid Type
Pleural Mesothelioma, Biphasic Type
Pheochromocytoma
Perihilar Cholangiocarcinoma
Paraganglioma
Papillary Thyroid Cancer
Papillary Stomach Adenocarcinoma
Papillary Renal Cell Carcinoma
Pancreatic Adenocarcinoma
Oligodendroglioma
Oligoastrocytoma
Myxofibrosarcoma
Mucinous Stomach Adenocarcinoma
Mucinous Carcinoma
Mucinous Adenocarcinoma Colon/Rectum
Mixed Germ Cell Tumor
Metaplastic Breast Cancer
Malignant Peripheral Nerve Sheath Tumor
Lung Squamous Cell Carcinoma
Lung adenocarcinoma
Low-Grade Glioma (NOS)
Leiomyosarcoma
Invasive Breast Carcinoma
Intrahepatic Cholangiocarcinoma
Intestinal Type Stomach Adenocarcinoma
Hepatocellular Carcinoma
Head and Neck Squamous Cell Carcinoma
Glioblastoma multiforme
Fibrolamellar Carcinoma
Esophageal Squamous Cell Carcinoma
Esophageal Adenocarcinoma
Endometrioid Carcinoma
Endocervical Adenocarcinoma
Embryonal Carcinoma
Diffuse Type Stomach Adenocarcinoma
Diffuse Large B-Cell Lymphoma, NOS
Desmoid/Aggressive Fibromatosis
Dedifferentiated Liposarcoma
Cutaneous Melanoma
Colon Adenocarcinoma
Chromophobe Renal Cell Carcinoma
Cholangiocarcinoma
Cervical Squamous Cell Carcinoma
Breast Invasive Mixed Mucinous Carcinoma
Breast Invasive Lobular Carcinoma
Breast Invasive Ductal Carcinoma
Breast Invasive Carcinoma (NOS)
Bladder Urothelial Carcinoma
Astrocytoma
Adrenocortical Carcinoma
D
MRGPRX4 promoter methylation (β)
p = 1.4x10-9
Nevus
Melanoma
E
Cell lines with damaging MRGPRX4 mutation
0/148
4/1807
Melanoma
Other cancers
F
Samples with damaging MRGPRX4 mutation
8/468
80/4660
Melanoma (SKCM)
all other TCGA cancers
G
NA
TCGA–UCEC
TCGA–LUAD
TCGA–ACC
TCGA–SKCM
TCGA–LUSC
TCGA–SARC
TCGA–COAD
TCGA–THYM
TCGA–HNSC
TCGA–STAD
TCGA–READ
TCGA–PRAD
TCGA–BRCA
TCGA–GBM
TCGA–CESC
TCGA–LIHC
TCGA–KIRC
TCGA–OV
TCGA–THCA
TCGA–LGG
Percent of samples with damaging MRGPRX4 mutation
H
Cutaneous_mel_patient_1
Cutaneous_mel_patient_2
MRGPRX4/Hoechst
I
melanoma states
Melanocytic
Neural crest-Like
Transitory
Undifferentiated
PC1
PC2
J
MRGPRX4
log(1+expression)
PC1
PC2
K
MRGPRX4 expression (Log2+1)
*
****
melanocytic
transitory
neural crest
undifferentiated

**Supplementary Figure 1. EMT/NC-Specific Upregulation of MRGPRX4 Expression in Melanoma Without Evidence of Recurrent Mutations**

(A-B) Normal human skin cell scRNA-seq. (A) UMAP of normal human structural skin cells (melanocytes, fibroblasts, keratinocytes and eccrine cells, GSE151091). (B) Expression of *MRGPRX4*, *TYR*, *KRT1*, *COL1A1* and *AQP5* in these clusters.

(C) Distribution of *MRGPRX4* mRNA expression across the detailed TCGA PanCancer Atlas

(D) *MRGPRX4* methylation in human nevus and cutaneous melanoma samples (GSE120878). β-value denotes the proportion of methylated CpG signal (0–1) across promoter-associated probes at the *MRGPRX4* locus.

(E-F) Damaging *MRGPRX4* mutations are rare. (E) *MRGPRX4* mutations in DepMap melanoma cell lines (0/148) or non-melanoma cancer lines (4/1807). (F) In TCGA, damaging *MRGPRX4* mutations appear in 8/468 SKCM samples, compared with 60/4000+ across all other cancers.

(G) Proportion of damaging *MRGPRX4* mutations across TCGA cancers

(H) Tile-scanned human cutaneous melanoma sections at 20x magnification, MRGPRX4 (red) expression localizes to discrete focal clusters of tumor cells.

(I,J) PCA of DepMap melanoma cell lines (I) PCA of DepMap melanoma cell lines stratified by established melanoma transcriptomic states (melanocytic, neural crest-like, transitory, undifferentiated).(J) PCA of the same samples overlaid with *MRGPRX4* mRNA expression

(K) Boxplot of *MRGPRX4* mRNA expression across melanoma states in melanoma cell lines (GSE80829). Data are analyzed by one-way ANOVA followed by post hoc test. $*p < 0.05$, $****p < 0.001$

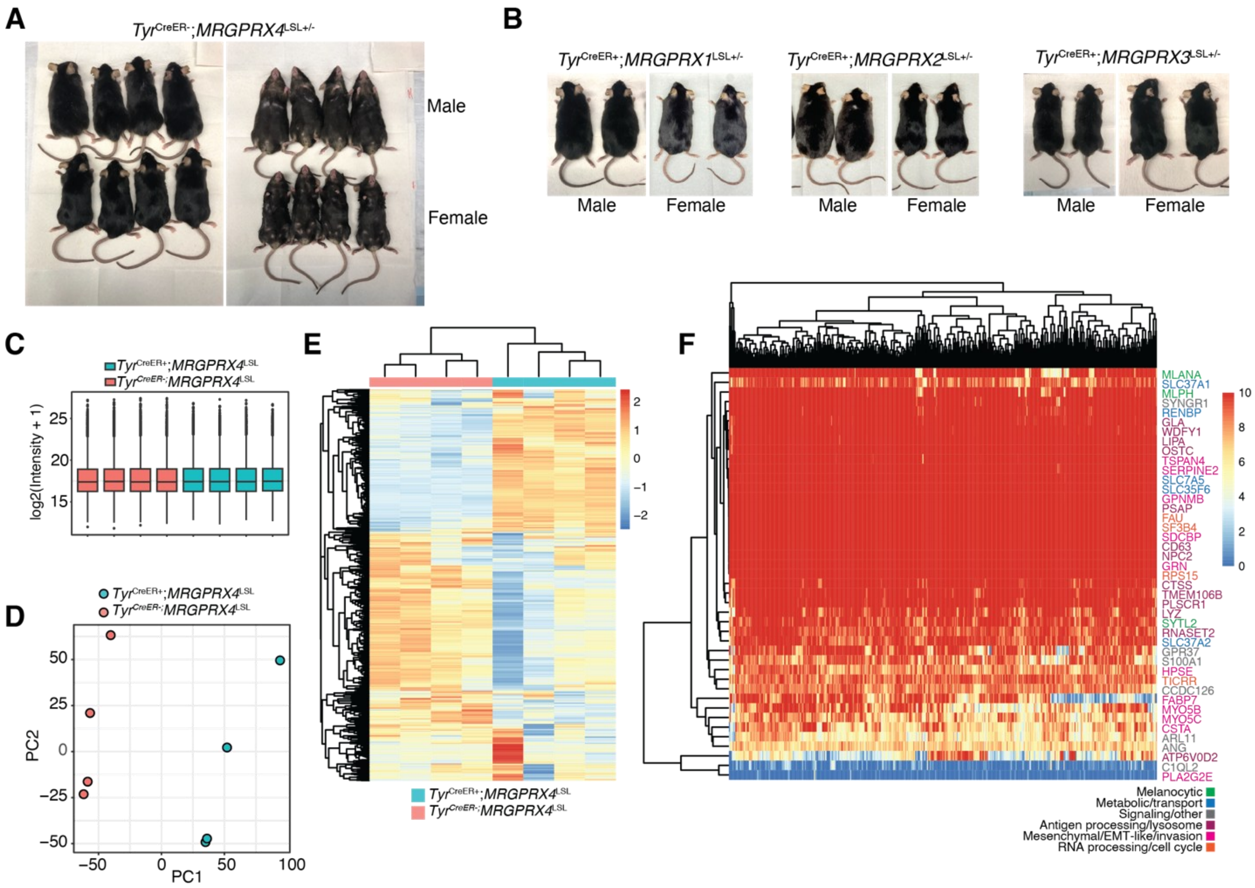


**Supplementary Figure 2. MRGPRX4 specifically drives melanoma in mouse.**

(A) *Tyr*$^{CreER-}$;*MRGPRX4*$^{LSL+/-}$ mice do not develop melanoma (analyzed up to 1.5 years)

(B) *Tyr*$^{CreER+}$;*MRGPRX1*$^{LSL+/-}$, *Tyr*$^{CreER+}$;*MRGPRX2*$^{LSL+/-}$, and *Tyr*$^{CreER+}$;*MRGPRX3*$^{LSL+/-}$ do not develop melanoma (analyzed up to 1.5 years)

(C-E) Proteomics of skin samples from skin tumor samples (*Tyr*$^{CreER+}$;*MRGPRX4*$^{LSL+/-}$ ) or control skin (*Tyr*$^{CreER-}$;*MRGPRX4*$^{LSL+/-}$)

(C) Boxplots show similar overall $log_2$-intensity distributions between melanoma samples and control skin

(D) PCA of tumor and control skin samples, indicating robust genotype-associated proteomic differences.

(E) Unsupervised hierarchical clustering of the top differentially expressed proteins separates cutaneous melanoma samples from melanoma and control skin

(F) Heatmap showing the mouse tumor proteome projected onto melanoma transcriptome derived from human TCGA SKCM. Mouse *Tyr*$^{CreER+}$;*MRGPRX4*$^{LSL+/-}$ tumors display a dominant mesenchymal/EMT-like–invasion and RNA-processing/cell-cycle program. Gene colors indicate assignment to established TCGA melanoma state categories.

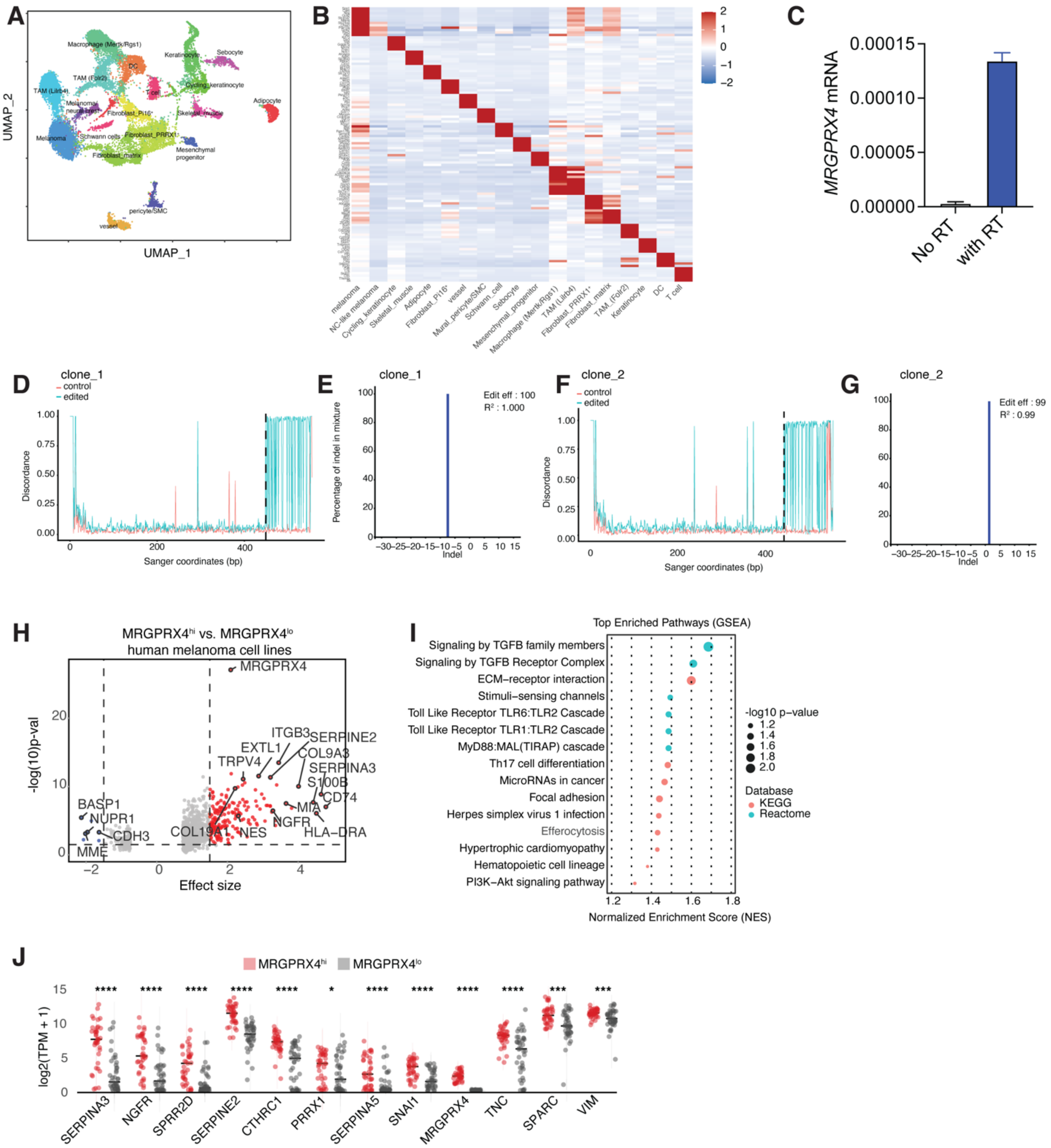


**Supplementary Figure 3. MRGPRX4 Defines a Mesenchymal and Invasive Melanoma Cell State**

(A,B) snRNAseq data from *Tyr*$^{CreER+}$;*MRGPRX4*$^{LSL}$ cutaneous melanoma tumors. (A) UMAP projection of all cell types within tumor samples showing major population clusters. (B) Heatmap of enriched genes for each cluster.

(C) *MRGPRX4* mRNA expression in A2058 melanoma cells

(D-G) Validation of *MRGPRX4* genomic disruption in A2058 cells for clone_1 and clone_2. (D) Clone_1 comparison of sequencing traces from edited and control cells showing a localized increase in discordance at the CRISPR cut position, consistent with loss of a sequence segment at the targeted site. (E) Analysis of insertions/deletions at this locus revealed a predominant −7 bp deletion accounting for ~100% of alleles, with indicated editing efficiency and model fit ($R^2$). (F) Clone_2 comparison of sequencing traces from edited and control cells showing a localized increase in discordance at the CRISPR cut position, consistent with loss of a sequence segment at the targeted site. (G) Analysis of insertions/deletions at this locus shows a predominant +2 bp deletion accounting for ~100% of alleles, with indicated editing efficiency and model fit ($R^2$). These frameshift mutations are predicted to introduce a premature stop codon and eliminate functional MRGPRX4.

(H-J) $MRGPRX4^{hi}$ melanoma cells display an ECM/EMT phenotype. (H) differential expression analysis of DepMap melanoma cell lines stratified as $MRGPRX4^{hi}$ or $MRGPRX4^{lo}$ based on endogenous transcript levels. $MRGPRX4^{hi}$ lines show increased expression of ECM-remodeling, EMT-associated, and inflammatory genes. (I) GSEA of ranked differential expression comparing $MRGPRX4^{hi}$ versus $MRGPRX4^{lo}$ DepMap melanoma lines. Top-enriched pathways include ECM-receptor interactions, TGFβ signaling, focal adhesion, innate sensing pathways, and PI3K-Akt signaling. (J) Expression of selected mesenchymal- and invasion-associated genes across $MRGPRX4^{hi}$ and $MRGPRX4^{lo}$ DepMap melanoma lines. ECM and EMT regulators are elevated in $MRGPRX4^{hi}$ lines.

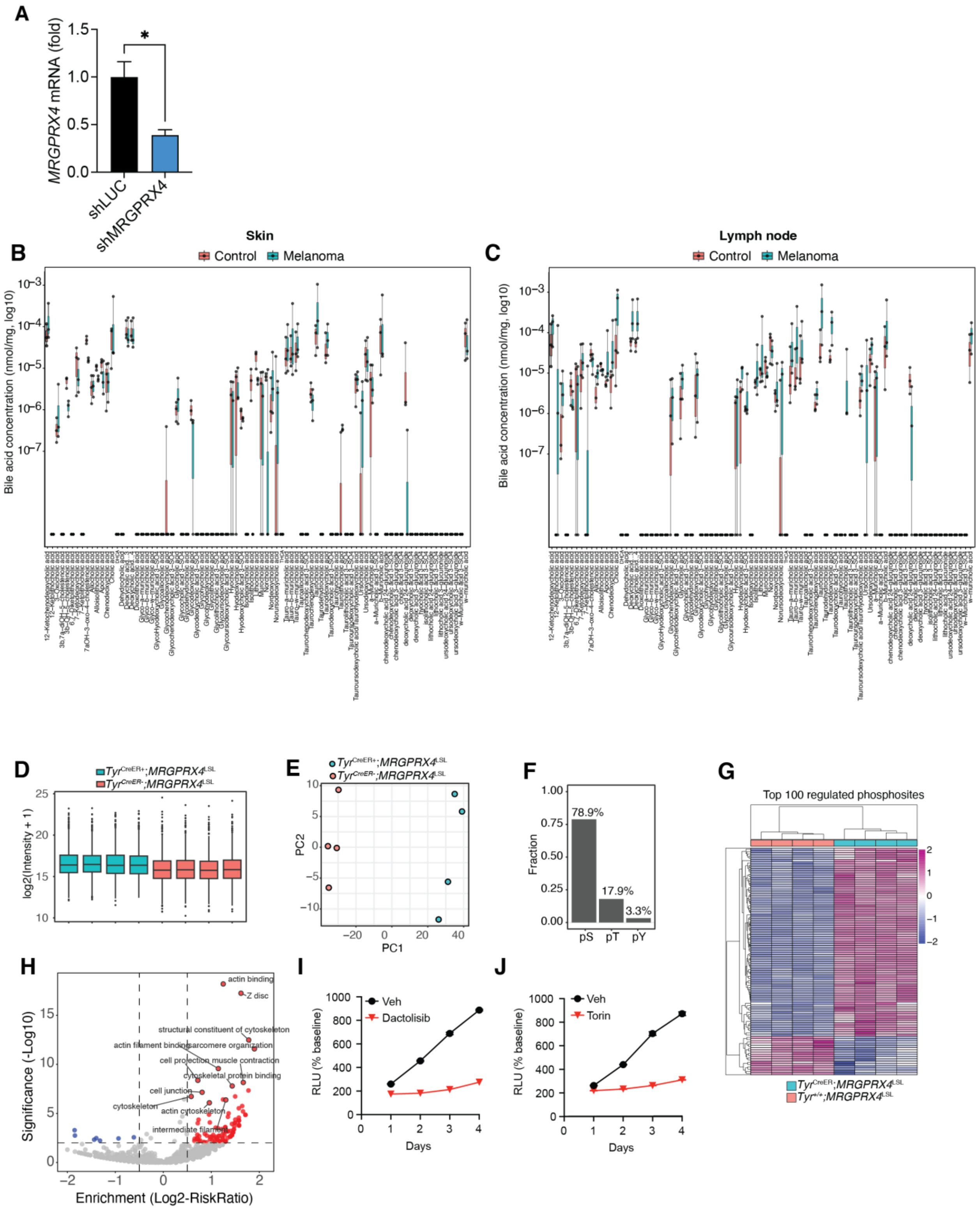


**Supplementary Figure 4. Additional validation of knockdown, lipidomics and phosphoproteomics**

(A) *MRGPRX4* mRNA expression in A2058 cells infected with lentivirus containing short hairpins targeting luciferase (shLuc) or MRGPRX4 (shMRGPRX4). Data is representative of 2-3 independent experiments

(B, C) Lipidomic quantification of bile acid species in (B) skin samples and (C) lymph nodes of skin tumor samples ($Tyr^{CreER+}$;$MRGPRX4^{LSL+/-}$ ) or control skin ($Tyr^{CreER-}$;$MRGPRX4^{LSL+/-}$)
(D-H) Phosphoproteomics of skin samples from skin tumor samples ($Tyr^{CreER+}$;$MRGPRX4^{LSL+/-}$ ) or control skin ($Tyr^{CreER-}$;$MRGPRX4^{LSL+/-}$). (D) Boxplots show similar overall $\log_2$-intensity distributions between tumor samples from $Tyr^{CreER+}$;$MRGPRX4^{LSL+/-}$ mice and control skin from $Tyr^{CreER-}$;$MRGPRX4^{LSL+/-}$ animals
(E) PCA of tumor and control skin, indicating robust genotype-associated proteomic differences.
(F) Abundance of phospho-residues in the phosphoproteomics dataset
(G) Unsupervised hierarchical clustering of the top differentially expressed proteins separates cutaneous melanoma samples from control skin.
(H) Volcano plot showing enriched phosphoproteomics pathways. Red dots indicate pathways enriched in tumor.
(I,J) A2058 cell proliferation, using ATP assay, in the presence of (I) 50 nM Dactolisib and (J) 50 nM torin. Data is representative of 2 independent experiments
Data are analyzed by Student's t test $^*p < 0.05$
